# Structure-Function Analysis of an Infantile Rhabdomyosarcoma Fusion-Oncogene Reveals Domain Requirements for Tumorigenesis

**DOI:** 10.64898/2026.11.01.763468

**Authors:** Chinmay S. Sankhe, Delia Calderon, Cenny Taslim, Melissa Sammons, Lindsay Ryan, Katherine M. Silvius, Amy C. Gross, Olivier Delattre, Joanna Cyrta, Ryan D. Roberts, Genevieve C. Kendall

## Abstract

Rhabdomyosarcoma is a pediatric soft-tissue sarcoma that reflects impaired skeletal muscle differentiation. Spindle cell/sclerosing rhabdomyosarcoma is a histological subtype found in children and adults. In children under two, spindle cell rhabdomyosarcoma is genetically driven by fusion-oncogenes of transcription factors or co-activators important for skeletal muscle lineage commitment and development. The most common fusion-oncogene in this group is VGLL2::NCOA2, a fusion between vestigial-like family member 2 (VGLL2) and nuclear receptor coactivator 2 (NCOA2). Children with fusion-driven rhabdomyosarcoma are treated with general chemotherapy, surgery, and radiation, which are not specific to the molecular underpinnings, and increase risk of secondary malignancies and premature death. Previously, we showed that VGLL2::NCOA2 is transforming in mouse myoblasts and transgenic zebrafish tumor models. Here, we build on this work by detailing the structure-function relationship of the retained VGLL2::NCOA2 protein domains to define domain requirements and their molecular interactions for tumorigenesis. We created deletion mutants of the VGLL2 vestigial/Tondu (TDU) domain and the NCOA2 activating domains 1 and 2 (AD1 and AD2), and a double mutant lacking the TDU+AD1 domains and assessed their function in a mouse myoblast model. We find that the TDU and AD1 domains support nuclear fusion-oncogene localization, transformation in vitro, and tumorigenesis in a mouse allograft model. Furthermore, transcriptomic analysis demonstrates that the TDU or AD1 mutant VGLL2::NCOA2 drives an upregulation of myogenesis gene sets and supports terminal differentiation. Overall, our results indicate that VGLL2::NCOA2 requires the TDU and AD1 domains for optimal transformation, and that targeting these domains, or their interacting protein partners, represents a therapeutic opportunity.

**Statement of Significance:** Understanding the contributions of the VGLL2::NCOA2 fusion-oncogene’s retained protein domains for tumorigenesis will refine potential molecular targets and identify new therapeutic opportunities in infantile rhabdomyosarcoma.

## Introduction

Sarcomas are predominantly genetically driven by gene fusion events that create aberrant transcription factors or kinases often with gain of function activities (1–4). This is true in infantile spindle cell/sclerosing rhabdomyosarcoma (RMS) that presents in children under the age of two (5–9). The most common fusion-oncogene in this infantile RMS subtype is the VGLL2::NCOA2 fusion-oncogene, which results from an inversion and translocation event that juxtaposes vestigial-like family member 2 (VGLL2) and nuclear receptor coactivator 2 (NCOA2). Additional fusion-oncogenes are also found in the disease, including VGLL2::CITED2, TEAD1::NCOA2, and SRF::NCOA2 (5–10). These tumors harbor few other mutations, suggesting that fusion-oncogenes are the founding and defining oncogenic drivers. Further, there are no therapies that directly target the genetics of the disease. In general, the tumors are responsive to standard of care and considered to have good prognosis; however, in some cases, they are also capable of metastasis and are challenging to treat if the tumor cannot be resected (7). The standard of care consists of general chemotherapy, which is non-specific and broadly toxic, with lifelong risk of secondary malignancy and premature death (11). Given that these fusion-oncogenes were identified within the past decade, the biological mechanisms underlying this disease are still being understood.

VGLL2, also known as VITO-1, was originally identified in a wingless Drosophila mutant (12) (also extensively reviewed in (13)). Further study in drosophila S2 cells found that TEAD facilitated the nuclear retention of vestigial protein and together, activated wing patterning and cell fate (14). In vertebrates, VGLL2 is specifically expressed during fetal stages in the somitic myotome and pharyngeal pouches and in adult skeletal muscle (15). In C2C12 mouse myoblasts, knocking down VGLL2 suppresses myogenic differentiation, whereas VGLL2 overexpression in chicken myoblasts promotes myogenic differentiation, including increasing chick myofiber diameter and density (16, 17). Vgll2-knockout mice have a redistribution of skeletal muscle fiber types to predominantly faster contractile muscle and a concurrent loss of exercise endurance (18). This is potentially due, in-part, to the loss of direct interaction between VGLL2 and TEAD1/TEAD4 transcription factors in neonatal skeletal muscles. VGLL2 and TEAD1 (also known as TEF-1) direct interaction requires the VGLL2 vestigial/TONDU (TDU) domain (17), and this binding allows VGLL2 to shuttle from the cytoplasm to the nucleus to promote skeletal muscle differentiation in cell culture (19). The nuclear function of VGLL2 is to re-direct TEAD binding at targets during skeletal muscle development and differentiation.

NCOA2 (also referred to TIF2, GRIP1, SRC2) is a member of the p160 steroid nuclear receptor coactivator family and regulates a range of processes, including, growth, differentiation, and metabolism (20, 21). NCOA2 is expressed in both proliferating myoblasts and differentiated myotubes and mediates the activation of myogenin (MyoG), MADS protein MEF-2, and regulator of cell cycle protein p21 (22). MyoG is a key regulator of skeletal muscle differentiation, and cells that express MyoG are locked in an intermediate differentiation state. The N-terminal of NCOA2 contains bHLH-PAS (basic helix loop helix-Per Arnt Sims) domain that also consists of the nuclear localization sequences (NLS) (23, 24). NCOA2 can facilitate histone acetylation and chromatin remodeling through the recruitment and direct binding of histone acetyltransferase p300/CBP to its activating domain 1 (AD1) (20, 25). The activating domain 2 (AD2) domain is responsible for interacting with histone methyltransferases of coactivator-associated arginine methyltransferase-1 and protein arginine methyltransferase 1 (CARM1/PRTM1) (26). Both AD1 and AD2 domains are located at the C-terminal of NCOA2 and are critical for regulation of transcriptional machinery in a normal context.

Previously, we functionally validated that VGLL2::NCOA2 is an oncogene that transforms in cross-species contexts including C2C12 mouse myoblasts and transgenic zebrafish. Further, when allografted into immunocompromised mice, C2C12 myoblasts expressing the VGLL2::NCOA2 fusion-oncogene develop aggressive tumors. Both the zebrafish and mouse allograft tumors histologically and transcriptionally recapitulate the human disease, and we leveraged these cross-species models to determine how VGLL2::NCOA2 engages developmental programs for tumorigenesis. This is mediated in-part through re-activation or inappropriate persistence of a small GTPase, ARF6 (27), which suggestively has multiple TEAD-transcription factor binding sites in the promoter region. Another group built on this work and showed that both VGLL2::NCOA2 and TEAD1::NCOA2 directly bind to TEAD transcription factors and p300 epigenetic factors, and engage TEAD-transcription in a 293T embryonic kidney cell model (28). Further, VGLL2::NCOA2 or TEAD1::NCOA2-driven allograft tumor models that were treated with systemic A485, a p300 small molecule inhibitor, had suppressed growth (28). These activities are likely related to the endogenous function of the VGLL2 and NCOA2 protein domains that are retained in the VGLL2::NCOA2 fusion-oncogene, but linking each protein domain’s functional impact on tumor formation has not been explored.

Here, we use a structure-function approach to understand how VGLL2::NCOA2 domains, and their broader molecular interactions, function independently and cooperatively to support fusion-driven oncogenesis. We used our C2C12 mouse myoblast system to test the contributions of the VGLL2 vestigial/TONDU (TDU) domain and NCOA2 activating domains 1 and 2 (AD1 and AD2) to tumorigenesis (17, 20, 22, 25, 26, 29). These domains are retained in the generation of the VGLL2::NCOA2 fusion-oncogene. We evaluated the VGLL2::NCOA2 fusion-oncogene and domain mutants’ cellular localization, transcriptional consequences, colony formation capacity, and in vivo transformation in mouse allografts. We found that the deletion of either the TDU or AD1 domains leads to a decrease in nuclear localization of the fusion-oncogene compared to full-length VGLL2::NCOA2 expressing cells. In addition, C2C12 myoblasts containing the VGLL2::NCOA2 TDU or AD1 domain deletion have a reduction in colony formation, delayed tumor onset in mouse allografts, and distinct histological features. RNA-sequencing analysis of the VGLL2::NCOA2 TDU or AD1 domain deletion mutants compared to full-length VGLL2::NCOA2 expressing myoblasts highlighted an upregulation in hallmark myogenesis gene sets and downregulation of cell cycle genes. These findings establish a critical role of the TDU and AD1 domains in VGLL2::NCOA2-driven transformation with a mechanism that is likely related to interfacing with co-factors that support an inhibition of skeletal muscle differentiation pathways.

## Materials and Methods

### Plasmids

We used our previously published VGLL2::NCOA2 full-length construct as the backbone for generating FLAG tagged and domain deletion constructs. This VGLL2::NCOA2 construct includes the coding sequence of *VGLL2* exons 1-2 linked to *NCOA2* exons 14-23 (27). We added a 3x FLAG tag (DYKDHD-G-DYKDHD-I-DYKDDDDK) to the 3’ of the full-length VGLL2::NCOA2 fusion-oncogene. We synthesized domain deletion mutants and point mutations based on previous literature and prediction models that indicated the location of the important domains within VGLL2 and NCOA2 (6, 19, 30–33). All experiments were performed with a FLAG tagged construct. These FLAG tagged and mutant constructs were synthesized by GenScript and then subcloned using the BamHI/NotI restrictions sites into the pcDNA3.1 (+) vector backbone (Invitrogen, #V79020) (27). Synthesized plasmids included the following deletions: TDU domain mutant deletion from amino acids 85 to 109, AD1 domain mutant deletion from amino acids 1010 to 1131, and the AD2 mutant deletion from amino acids 1288 to 1464. Plasmid DNA sequences were confirmed with whole plasmid sequencing using Plasmidsaurus.

### Cell culture

C2C12 mouse myoblast cells were obtained from American Type Culture Collection (ATCC Cat# CRL-1772, RRID:CVCL_0188) and were maintained in a humidified incubator at 37^°^C and 5% CO_2_. Before transfection, C2C12 cells were cultured in growth media that included Dulbecco’s Modified Eagle Medium (DMEM) with GlutaMAX supplement (Gibco #10-569-044), 10% Fetal Bovine Serum (Corning #35016CV), and 1x Penicillin-Streptomycin (Pen/Strep) (Gibco #15-140-122). After transfection, C2C12 cells were cultured in growth media that included Dulbecco’s Modified Eagle Medium (DMEM) with GlutaMAX supplement (Gibco #10-569-044), 10% Fetal Bovine Serum (Corning #35016CV), and 1 mg/ml Geneticin/G418 antibiotic (Gibco #10131027). Cells were tested for mycoplasma and were confirmed negative by qPCR (Genetica, Lab Corp), and cell identity was verified by STR profiling routinely (at least every six months).

### C2C12 cell transfection

C2C12 cells were transfected using the GenJet DNA InVitro Transfection Reagent (SignaGen Laboratories #SL100489) using the manufacturer’s recommended protocol. The control group was transfected with the pcDNA3.1 (+) empty vector, and the experimental groups were transfected with the pcDNA3.1-VGLL2::NCOA2 construct or the pcDNA3.1-VGLL2::NCOA2 mutant domain deleted constructs. All constructs were tagged with 3x FLAG at the 3’ end. Transfected cells were maintained in growth media without the addition of Pen/Strep along with selection using 1 mg/ml Geneticin/G418 antibiotic (Gibco #10131027) for a minimum of two weeks prior to all experiments, and were maintained in this media.

### Immunofluorescence staining

Transfected C2C12 cells were seeded on 22-mm diameter circular collagen-coated coverslips (Corning® BioCoat® Collagen I #354089) placed in a 6-well plate at a density of 50,000 cells per well. Cells were allowed to adhere for 48 hours. Cell samples were fixed using 4% paraformaldehyde in 1x phosphate buffered saline (PBS) for 15 min at room temperature. The samples were rinsed with 1x PBS twice and then incubated with permeabilization solution consisting of 0.25% Triton X-100 in 1x PBS for 10 min at room temperature. The permeabilization solution was aspirated, and samples were incubated in blocking buffer consisting of 5% goat serum and 0.1% Triton X-100 in 1x PBS for 1 hour at room temperature. The samples were then incubated overnight at 4°C with the primary antibody solution consisting of a FLAG antibody (FLAG primary antibody used at 1:500, Sigma-Aldrich #F1804, RRID:AB_262044) diluted in 3% BSA in 1x PBS and 0.1% Tween (PBST). The samples were rinsed three times with 1x PBS on a gentle rocker for 5 min and then incubated for 1 hour at room temperature with secondary antibody (goat anti-mouse Alexa Fluor 488 secondary antibody used at 1:500, Invitrogen #A32723) diluted in blocking buffer. The samples were again rinsed three times with 1x PBS and mounted on a microscope slide using DAPI-containing mounting medium for counterstaining nuclei of the cells (ProLong™ Gold Antifade Mountant with DNA Stain DAPI, Invitrogen #P36935). Immunofluorescence staining experiments were repeated three times for the cells transfected with pcDNA3.1 control, full-length VGLL2::NCOA2, and mutants with deletions of TDU, AD1, AD2, AD1+AD2 double mutant, and TDU+AD1 double mutant.

### Immunofluorescence microscopy and imaging analysis

Immunofluorescence staining samples were imaged using a Keyence BZ-X810 fluorescence microscope on a PlanFluor 40x 0.60/3.30-2.20mm Ph2 objective lens and a high-resolution monochrome camera setting. The exposure time for capturing the FLAG stain was kept constant (3 sec) for imaging all the samples. The correlation maps between the nuclear stain (DAPI) and the FLAG stain were generated using the Keyence analyzer software. Images (DAPI and FLAG channels) were converted to 16-bit and were analyzed simultaneously using JaCoP colocalization Plugin (34) (RRID:SCR_025164) on FIJI (RRID:SCR_002285) to evaluate the overlap coefficient. If the overlap coefficient is higher, there is an increase in the co-localization of the DAPI and the FLAG stain positive pixels, indicating higher correlation.

### Western blotting

For extraction of whole-cell protein lysates, cells were lysed in 150 µl of RIPA buffer (Sigma-Aldrich #R0278) supplemented with 10 µl per ml of the 100x Halt protease and phosphatase inhibitor cocktail (Thermo Scientific #78442) and briefly homogenized using a pellet pestle. The lysates were incubated on ice for 45 min with periodic vortexing every 5-10 min. The cell lysates were centrifuged at max speed at 4°C for 15 min and the supernatant was transferred into a new tube. The lysate concentration was quantified using the Pierce BCA Protein Assay Kit (Thermo Scientific #23227). Protein lysates were denatured using 2X Laemmli buffer (Bio-Rad #1610737) at 95°C for 10 min before electrophoresis.

For extraction of the nuclear and the cytosolic protein fractions, cells were lysed in 75 µl of mild cell lysis buffer containing 1 M Tris-HCl, 10 mM NaCl, 10% NP-40 and Halt Protease and Phosphatase inhibitor cocktail. Lysates were homogenized using a pellet pestle and rotated at 4°C for 10 min. The solution was centrifuged at 1300g at 4°C for 5 min, and the supernatant (cytosolic fraction) was transferred into a new tube. The pellet was washed with 50 µl of mild cell lysis buffer and centrifuged at 4°C for 5 min at 1300g. The supernatant was discarded, and the pellet (nuclear fraction) was resuspended in 24 µl of lysis buffer, and the concentrations of both the cytosolic and nuclear fractions were quantified using the Pierce BCA Protein Assay Kit.

Protein lysates were denatured using 2x Laemmli buffer with boiling at 95°C for 10 min prior to electrophoresis. The denatured protein lysates were separated on 4-15% Mini-PROTEAN® TGX Precast Protein Gel with 10 ul of Precision Plus Protein™ Dual Xtra Prestained Protein Standard (Bio-Rad #1610377) as a ladder at 100 V for 90 min at room temperature. The proteins on the gel were transferred on a Immun-Blot PVDF membrane (Bio-Rad #1620177) using a wet transfer setup and blocked with Casein (Thermo Scientific #PI37528) + 0.05% Tween for 60 min, followed by incubation with FLAG primary antibody (1:1000, Sigma-Aldrich #F1804, RRID:AB_262044) overnight on an orbital shaker at 4°C. The membrane was washed with 1x phosphate buffered saline + 0.1% v/v Tween (PBST) three times on a shaker for 10 min and then incubated with secondary antibody (1:10000; goat anti-mouse IgG (H + L)-HRP Conjugate, Bio-Rad #1706516) in Casein + 0.05% Tween for 1 hour at room temperature. The membrane was washed with 1x PBST three times for 10 min with agitation at room temperature before proceeding to visualization. For visualization, we used SuperSignal^TM^ West Atto Ultimate Sensitivity Substrate (Thermo Scientific #PI34577) and imaged on the C-DiGit Chemiluminescent LI-COR imager or ChemiDoc Go Imaging System (Bio-Rad). For probing of loading control, the membrane was stripped for 20 min with the Thermo Scientific™ Restore™ Western Blot Stripping Buffer (ThermoScientific #PI21059) and washed with 1x PBST three times for 5 min on a rocker at room temperature. The membrane was blocked with Casein + 0.05% Tween for 30 min, followed by incubation with Tbp (1:1000, Cell Signaling #44059, RRID:AB_2799258) or β-actin (1:1000, Invitrogen #MA532540, RRID:AB_2809817) primary antibody overnight with agitation at 4°C. The membrane was washed as described above and incubated with secondary antibody (1:10000; goat anti-rabbit IgG (H + L)-HRP Conjugate, Bio-Rad #1721019) for 1 hour at room temperature. Washing and imaging of the membrane was performed as described above. Western blots were quantified by densitometry analysis using ImageJ (RRID:SCR_003070) or FIJI.

### Soft agar colony formation assay

Colony formation assays were performed in 6-well plates using an adapted protocol (35, 36). Briefly, 1% noble agar (Research Products International #A20460) was mixed with 2x cell culture medium (DMEM, Fisher #12-800-017; and 0.2g sodium bicarbonate dissolved in nuclease-free water) and left to solidify for 30 minutes. 0.6% noble agar mixed with cell culture medium containing 1mg/mL of Geneticin/G418 was used as the top layer. The top layer consisted of 80,000 cells per plate. We included 100uL of cell culture medium with 1 mg/ml Geneticin/G418 to avoid dryness of the top layer. We added 100-200uL of fresh medium bi-weekly to the top layer for 2 weeks. After 2 weeks, 200uL of 1mg/mL of NitroBlue Tetrazolium Chloride (Fisher #N6495) in 1x PBS was added to each well for 24 hours at 37°C. The plates were then imaged using Bio-Rad gel imager or ChemiDoc Go Imaging System (Bio-Rad) and colonies were counted manually using ImageJ or FIJI.

### Mouse allograft experiments and tumor processing

C2C12 cells transfected with the full-length VGLL2::NCOA2 fusion-oncogene and the mutant fusions having deletions of TDU, AD1, or TDU+AD1 domains were evaluated in mouse allograft models. C2C12 cells were transfected and selected with Geneticin/G418 for at least two weeks prior to mouse allograft experiments. Two million cells were resuspended in 20 µl of sterile 1X PBS and were injected intramuscularly/orthotopically into the gastrocnemius muscle of one leg of a severe combined immunodeficient (Icr--SCID, C.B-17/IcrHan Hsd-Prkdc^scid^) mouse under anesthesia. The groups contained male mice between 36-42 days of age. There were 8 mice per injection group for Trial 1 (total of 32 mice) and 6 mice per injection group for Trial 2 (total of 24 mice). The mice were monitored for tumor mass growth, and after tumor detection, the volume of the tumor was measured biweekly using calipers until the tumor burden reached endpoint (volume of 1000 mm^3^) or until the end of the experiment (122 days post injection for Replicate 1, 100 days post injection for Replicate 2). There were 2 mice (1 in AD1 domain deletion group and 1 in TDU+AD1 domain deletion group) in Replicate 1 and 1 mouse (in TDU+AD1 domain deletion group) in Replicate 2 that had to be euthanized for humane considerations prior to any tumor formation, and no tissues were collected from these mice. Tumors were fixed in 10% neutral buffered formalin for 24 hours, then stored in 1xPBS until processing.

### Histopathology analysis

Tumors were mounted in paraffin blocks for microtome sectioning, and H&E stains were performed at the Nationwide Children’s Hospital (NCH) Histopathology, Processing, and Banking Core. All slides were scanned on a Hamamatsu Nanozoomer S360 scanner (Hamamatsu, Japan) with a 40x objective. Morphology of tumor cells was assessed on hematoxylin and eosin (H&E)-stained slides by a pathologist blinded to the experimental condition, in a semi-quantitative manner, using a three-tiered system: mostly (>75%) round/pleomorphic cells, mostly (>75%) spindle cells, or a combination of the above. Quantification of tumor and necrosis areas on H&E slides, was performed using the HALO image analysis platform and the HALO AI module (version 3.5.3577) from Indica Labs (Albuquerque, NM, USA). Outputs for all slides were visually inspected and validated by a pathologist. Percentage necrosis was evaluated by dividing the area of necrosis to the total allograft section area (tumor area + necrosis area) and multiplying by 100.

### Immunohistochemistry

Automated myogenin immunohistochemistry was performed using a Bond III immunostainer (Leica Microsystems, Nanterre, France), the Bond Polymer Refine detection system, and the anti-myogenin antibody clone F5D (Dako, Stockholm, Sweden) (pH9 retrieval, 1:100 dilution, 60 min incubation). All slides were scanned on a Hamamatsu Nanozoomer S360 scanner (Hamamatsu, Japan) using 40x objective. Quantification of the percentage of myogenin-positive tumor nuclei on stained slides was performed using the HALO image analysis platform and the HALO AI module (version 3.5.3577) from Indica Labs (Albuquerque, NM, USA).

Manual immunohistochemistry was performed for myogenin and myosin heavy chain (MF20) on the mouse allograft tumor sections. For myogenin (MyoG) and myosin heavy chain immunohistochemistry, slides were baked in a hybridization oven at 60°C for 1 hour followed by deparaffinization with Histo-Clear (National Diagnostics #HS2021GLL) twice for 10 min each. The slides were rehydrated with two washes of 100% ethanol for 3 min each, two washes of 95% ethanol for 3 min each, and RO water for 3 min. Antigen retrieval was performed in 10 mM sodium citrate buffer pH 6.0 for 20 min in a pressure cooker, maintaining the temperature of the buffer to be at least 95°C. The slides were cooled for 5-10 min and washed with 1x PBS twice for 3 min each. Blocking was performed for 1 hour in a humid chamber at room temperature using 3%(w/v) bovine serum albumin (BSA) in 1x PBST with 10%(v/v) of goat serum. The tumor sections were incubated with the primary antibody diluted in 3% BSA in 1x PBST and incubated in a humid chamber at 4°C overnight. The primary antibodies included myogenin (1:100, F5D, DSHB, RRID:AB_528355) and MF20 (1:100, DSHB, RRID:AB_2147781). The slides were washed in 1x PBST four times on a rocker for 5 min each and incubated with 3% hydrogen peroxide solution for 30 min in a humid chamber. After washing with 1x PBS twice for 3 min each, the slides were incubated in secondary antibody solution (1:300 goat anti-mouse IgG (H + L)-HRP Conjugate (Bio-Rad #1706516) in 3% BSA in 1x PBST) for 1 hour at room temperature. Slides were washed in 1xPBST as described before and 3,3′-Diaminobenzidine tetrahydrochloride (DAB) (Sigma Aldrich #D5905) solution (1 ml) containing 1 µl of 30% hydrogen peroxide (Sigma-Aldrich) was applied to the sections and incubated for 5-7 min with periodic evaluation to determine the development of stain. DAB solution was rinsed off using RO water.

For FLAG immunohistochemistry on mouse allograft tumor sections, antigen retrieval was performed using 1x citrate unmasking solution (Cell Signaling #14746) and microwaving the slides for 10 min at a sub-boiling temperature (95-98°C). Slides were cooled for 30 min at room temperature, and incubated with 3% hydrogen peroxide solution for 10 min. Slides were blocked using 5%(v/v) goat serum in 1x TBST (Cell Signaling #9997) at room temperature for 1 hour. The primary FLAG antibody (1:200, Cell Signaling Cat #14793, RRID:AB_2572291) was diluted in SignalStain Antibody diluent (Cell Signaling #8112) and applied on the slides for overnight incubation. Then, the slides were washed in 1x TBST three times for 5 min each and incubated with SignalStain Boost detection reagent (Cell Signaling #8114) for 30 min at room temperature. For development of stains, slides were incubated with SignalStain DAB Chromagen concentrate diluted in SignalStain DAB diluent (Cell Signaling #8059) for 7-10 minutes. The slides were counterstained using hematoxylin (RICCA Chemical Company #3535-16) for 5 min and rinsed off with RO water for another 5 min. Samples were dehydrated in two incubations of 95% ethanol, 100% ethanol, and Histo-Clear for 1 min each. Mounting was performed using Prolong Gold Antifade Mountant (Invitrogen #P36934) and dried for 24 hours before sealing and imaging on a Keyence BZ-X810 microscope using a PlanFluor 40x 0.60/3.30-2.20mm Ph2 objective lens and a high-resolution color camera setting. The exposure time was 1/10 sec and was consistent for all the samples.

### RNA sequencing (RNA-seq) and analysis

Total RNA from transfected C2C12 cells was isolated using the RNeasy Mini Kit (Qiagen #74104) with on-column DNase digestion. A total of four replicates per transfection condition was sent for RNA-sequencing. RNA concentration and integrity were quantified using nanodrop spectrophotometer and Agilent TapeStation RNA ScreenTape and sample QC and library preparation were performed at the Institute for Genomic Medicine (IGM) at Nationwide Children’s Hospital. Libraries were run on a NovaSeq6000 S1 PE150bp paired-end reads, and a total of 60-80 million reads were generated as output per sample. RNA-seq analysis was performed using an in-house pipeline (RNA-SeqEZPZ) on the Nationwide Children’s Hospital high-performance cluster (37). Briefly, FASTQC (38) were used to check the quality of reads. Low-quality bases and adapter sequences were trimmed using Trim Galore (39). Reads were aligned to a custom mouse genome (mm39) that also included the human VGLL2::NCOA2 fusion-oncogene or the VGLL2::NCOA2 junction sequence using STAR (40) and were assigned to genomic features using featureCounts (41). DESeq2 (42) and SARTools (43) were used to perform differential gene analysis. Pathway analysis was performed using clusterProfiler (44) and the msigdb (45) database. Overlap analysis was done using GeneOverlap (46).

### Statistical analysis

All sample sizes and experimental replicates are detailed in the figure or the figure legends. The statistical analysis was performed using GraphPad Prism version 10.3.1 for Windows (RRID:SCR_002798, GraphPad Software, Boston, Massachusetts USA, www.graphpad.com). Normality was tested using the Shapiro-Wilk normality test performed using GraphPad Prism. The information on the statistical test performed on the respective experimental figure panel is indicated in the figure legend.

## Data Availability Statement

The RNA-seq data generated in this publication derive from mouse samples and have been deposited in NCBI’s Gene Expression Omnibus (47) and are accessible through GEO Series accession number GSE327933 (https://www.ncbi.nlm.nih.gov/geo/query/acc.cgi?acc=GSE327933)

The plasmids generated in this study are available through Addgene with the following identification numbers: TDU domain deletion - #242455, AD1 domain deletion - #242457, AD2 domain deletion - #255333, AD1+AD2 domain deletion - 255334, TDU+AD1 domain deletion - #242476, TDU alanine-rich mutation - #255335, AD1 alanine-rich mutation - #255336, AD1 point mutations - #255337.

The other reagents or data generated in this study are available upon request from the corresponding author.

## Results

### VGLL2::NCOA2 TDU and AD1 domains regulate subcellular localization

First, we investigated the independent roles of the TDU, AD1, and AD2 structural domains in supporting VGLL2::NCOA2 activity in cell culture. We used our C2C12 mouse myoblast model where the human form of the VGLL2::NCOA2 fusion-oncogene was transfected into C2C12 cells using a pcDNA3.1 (+) backbone. This model readily transforms in cell culture and in mouse allograft systems (27). We modified this approach by adding a 3x FLAG tag at the 5’ or 3’ end of VGLL2::NCOA2 to support detection and biochemical experiments. We tested the capacity of both FLAG tagged versions to transform C2C12 cells using colony formation and found that the 3’ FLAG tag had increased fusion-oncogene transformation efficiency compared to the 5’ FLAG tag (Figure S1A). Therefore, we proceeded with a 3’ FLAG tag for the full-length VGLL2::NCOA2 fusion-oncogene in all the following experiments. We also created 3’ FLAG domain deletion mutants of the VGLL2::NCOA2 fusion-oncogene by removing the TDU, AD1, or AD2 domains (Figures 1A, S1B). The VGLL2::NCOA2 or domain deletion mutant constructs were transfected into C2C12 mouse myoblasts and selected for two weeks with G418. We then performed a western blot and immunofluorescence staining to determine fusion-oncogene expression levels and to delineate subcellular localization.

**Figure 1:**
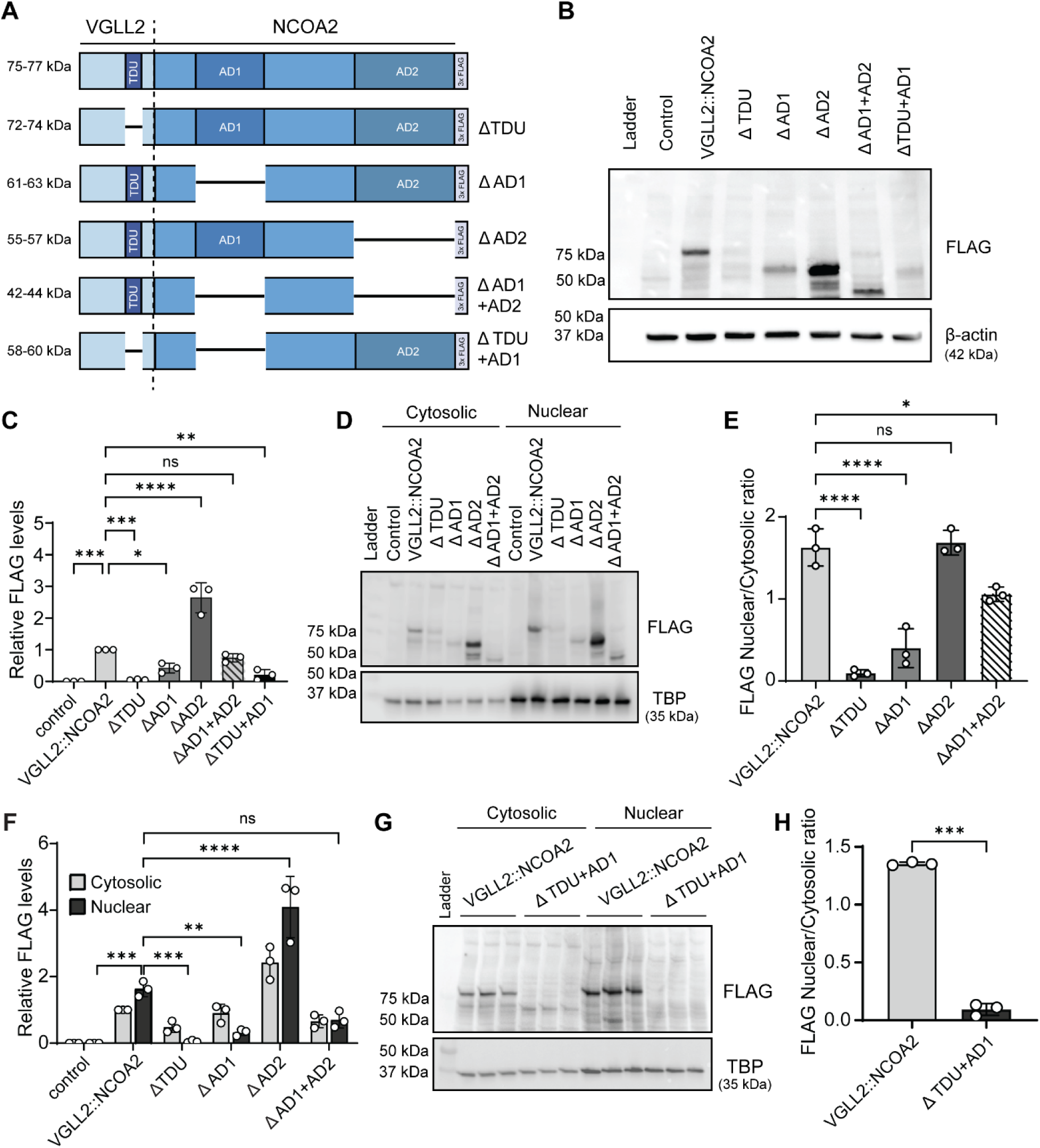
VGLL2::NCOA2 TDU and AD1 structural domains dictate subcellular localization. (A) Schematic depicting the structural domains of VGLL2::NCOA2 and regions removed in domain deletion mutants. C2C12 mouse myoblast cells transfected with pcDNA3.1 empty vector control or pcDNA3.1 expressing 3’ end 3x FLAG-tag VGLL2::NCOA2 and mutants having either: 1) Tondu/Vg (TDU) domain removed, 2) activating domain 1 or 2 (AD1 or AD2) removed, or 3) dual deletions of TDU+AD1 and AD1+AD2 domains. The predicted molecular weight of each construct is indicated to the left. (B) Western blot of whole-cell protein lysates extracted from transfected C2C12 cells that were selected with Geneticin/G418 for two weeks and probed for FLAG and loading control β-actin. The predicted molecular weights for the full-length fusion and deletion mutants are: full-length (∼75-77 kDa), TDU (∼72-74 kDa), AD1 (∼61-63 kDa), AD2 (∼55-57 kDa), AD1+AD2 (∼42-44 kDa), TDU+AD1 (∼58-60 kDa). (C) Quantification of relative levels of FLAG in C2C12 transfected cells via densitometry analysis from western blotting in Panel (B). Data is normalized with respect to the full-length VGLL2::NCOA2 condition. Data indicate mean ± SD for n=3 trials. The p values were calculated using a one-way ANOVA followed by Tukey’s multiple comparisons post hoc test. *p<0.05, **p<0.01, ***p<0.001, ****p<0.0001. (D) Western blot of cytosolic and nuclear protein lysates extracted from transfected C2C12 cells that were selected for two weeks and probed for FLAG and loading control Tbp. (E) Quantification of nuclear:cytosolic ratio of FLAG in C2C12 transfected cells via densitometry analysis from the western blot in Panel (D). Data indicate mean ± SD for n=3 trials. The p values were calculated using a one-way ANOVA followed by Tukey’s multiple comparisons post hoc test. *p<0.05, **p<0.01, ****p<0.0001, ns: not significant. (F) Quantification of relative nuclear and cytosolic FLAG levels in C2C12 transfected cells via densitometry analysis from the western blot in Panel (D). Data is normalized to the cytosolic full-length VGLL2::NCOA2 transfected condition. Data indicate mean ± SD for n=3 trials. The p values were calculated using a one-way ANOVA followed by Tukey’s multiple comparisons post hoc test. **p<0.01, ***p<0.001, ****p<0.0001, ns: not significant. (G) Western blot of three technical replicates of cytosolic and nuclear protein lysates extracted from C2C12 cells transfected with full-length VGLL2::NCOA2 and dual deleted TDU+AD1 domains and probed for FLAG and loading control Tbp. (H) Quantification of nuclear:cytosolic ratio of FLAG in C2C12 transfected cells via densitometry analysis from the western blot in Panel (G). Data indicate mean ± SD for n=3 technical replicates. The p values were calculated using a T-test with Welch’s correction. ***p<0.001.

Western blot from whole-cell protein lysates with the FLAG antibody showed that the fusion-oncogene was highly expressed in the VGLL2::NCOA2 full-length condition and not detected in the pcDNA control cells (Figure 1B). We observed variable protein expression of the fusion-oncogene in the mutant fusion transfected cells (Figure 1B). Compared to full-length VGLL2::NCOA2, there were higher levels of the AD2 mutant, and lower levels of the TDU, AD1, and TDU+AD1 mutants (Figure 1B,C). We determined that the oncogenic activity did not directly correlate to protein levels by using complementary mutations, including alanine substitutions in the TDU and AD1 domains, and key point mutations in the AD1 domain (Figure S2). After transfecting these fusion-oncogene mutants into C2C12 cells, there were similar fusion-oncogene protein levels compared to full-length VGLL2::NCOA2 (Figure S3). Transformation capacity was comparable between these TDU and AD1 alanine substitution and point mutations and the domain deletion mutants. All forms of the TDU and AD1 mutations were unable to transform myoblasts in cell culture (Figure S2, S3).

VGLL2 and NCOA2 are transcriptional coactivators, and the fusion of these two proteins drives transcriptional changes (27). Therefore, we hypothesized that the full-length fusion-oncogene translocates to the nucleus for full functionality, akin to VGLL2’s known activity during myogenesis (15–18). To test this, we probed for the subcellular localization of VGLL2::NCOA2 in C2C12 cells and studied the changes in localization following domain deletions. We isolated cytosolic and nuclear protein lysates and performed western blots and found that the full-length VGLL2::NCOA2 and single AD2 domain deletion mutant are primarily nuclear (Figure 1D,E). Deletion of the TDU and AD1 domains led to a reduction in nuclear localization, and compared to full-length VGLL2::NCOA2, an overall decrease in cytosolic and nuclear levels (Figure 1D-F). Dual AD1+AD2 domain deletion caused the fusion-oncogene to be pancellular (Figure 1D-F), while dual TDU+AD1 domain deletions retained fusion-oncogene localization in the cytosol with decreased cytosolic and nuclear levels compared to the full-length transfection condition (Figure 1G,H, S4A).

We also performed immunofluorescence staining for the full-length VGLL2::NCOA2 and mutants as a complementary approach for assessing localization (Figure 2A). FLAG staining on transfected myoblasts revealed that cells with the full-length VGLL2::NCOA2 and AD2 domain deletion mutant had more FLAG overlap with the nucleus compared to the control pcDNA cells. This was quantified by plotting the correlation maps of the DAPI (nuclear) stain for each pixel as compared to FLAG (fusion-oncogene) (Figure 2A,B). On the other hand, single deletion of the TDU and AD1 domains decreased the overlap of FLAG with the nuclear stain, suggesting reduced fusion-oncogene localization in the nucleus. Dual deletions of AD1+AD2 and TDU+AD1 also decreased nuclear localization of the fusion-oncogene (Figure 2A,B, S4B). Overall, our results indicate that the VGLL2 TDU and NCOA2 AD1 structural protein domains play a critical role in regulating subcellular localization of the VGLL2::NCOA2 fusion-oncogene, while the NCOA2 AD2 domain is not required for nuclear localization.

**Figure 2:**
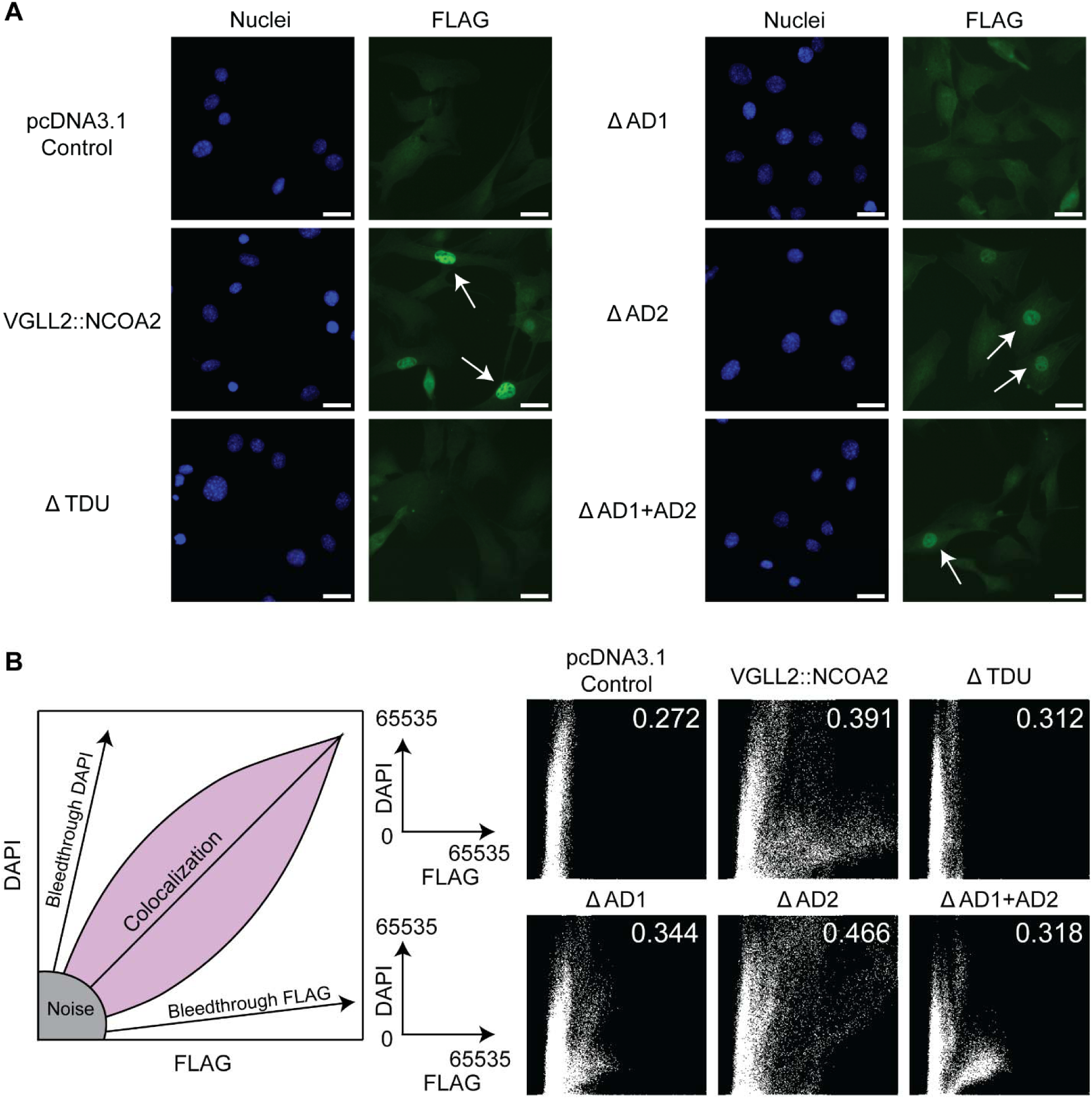
VGLL2::NCOA2 TDU and AD1 domains regulate nuclear translocation. (A) Immunofluorescence staining of FLAG in C2C12 cells transfected with the full-length VGLL2::NCOA2 fusion-oncogene and domain deletion mutants. White arrows indicate cells with nuclear localized FLAG (VGLL2::NCOA2). Scale bars: 25 µm. (B) Correlation maps depicting co-localization of DAPI nuclear stain with FLAG (VGLL2::NCOA2) for different transfected C2C12 conditions. The Y-axis depicts DAPI pixel intensity and the X-axis depicts FLAG pixel intensity. Each white point represents the pixel intensity for DAPI and FLAG at the same location on the image. The number (white, top right) on the colocalization maps indicates the overlap coefficient between FLAG and DAPI. The greater the number of white dots in the purple region (shown in the schematic) corresponds to higher colocalization/higher overlap coefficient between DAPI and FLAG.. These results are representative of n=3 trials.

### VGLL2::NCOA2 transformation capacity in cell culture is dependent on the TDU and AD1 domains

We next evaluated the functional consequences of removing these protein domains and how that influenced VGLL2::NCOA2 tumorigenic potential. Previously, we showed that C2C12 cells transfected with the full-length VGLL2::NCOA2 fusion-oncogene generated colonies in soft agar assays (27). Here, we plated C2C12 mouse myoblasts containing either full-length VGLL2::NCOA2 or domain deletion mutants in soft agar and monitored colony formation. After two weeks, we counted the colonies and found that the full-length VGLL2::NCOA2 construct and AD2 domain deletion construct resulted in on average ∼200-300 colonies per well (Figure 3A-D). Single TDU and AD1 domain deletions significantly suppressed colony formation compared to the full-length VGLL2::NCOA2 condition. Dual deletions of AD1+AD2 and TDU+AD1 also resulted in a significant reduction in colony number (Figure 3A-D). To validate that oncogenic potential was not dictated by protein expression level, we generated a TDU domain mutant with an alanine-rich sequence and transfected and selected the C2C12 cells for two weeks. Western blotting revealed similar protein levels of the mutant and full-length VGLL2::NCOA2 fusion-oncogene in transfected cells (Figure S2, S3A,C). We then performed a colony formation assay and found that similar to the TDU domain deletion mutant, the TDU-alanine rich mutant did not form colonies (Figure S3D,E). We also generated alanine-rich and multiple point mutations in the AD1 domain and achieved comparable protein expression to the full-length VGLL2::NCOA2 fusion-oncogene (Figure S2, S3B,C). Colony formation assay with these mutants was similar to that seen with the AD1 deletion mutant, with few colonies forming (Figure S3D,E). Overall, our results suggest that in this context the TDU and AD1 domains support the oncogenic potential of the VGLL2::NCOA2 fusion-oncogene, while the AD2 domain is dispensable.

**Figure 3:**
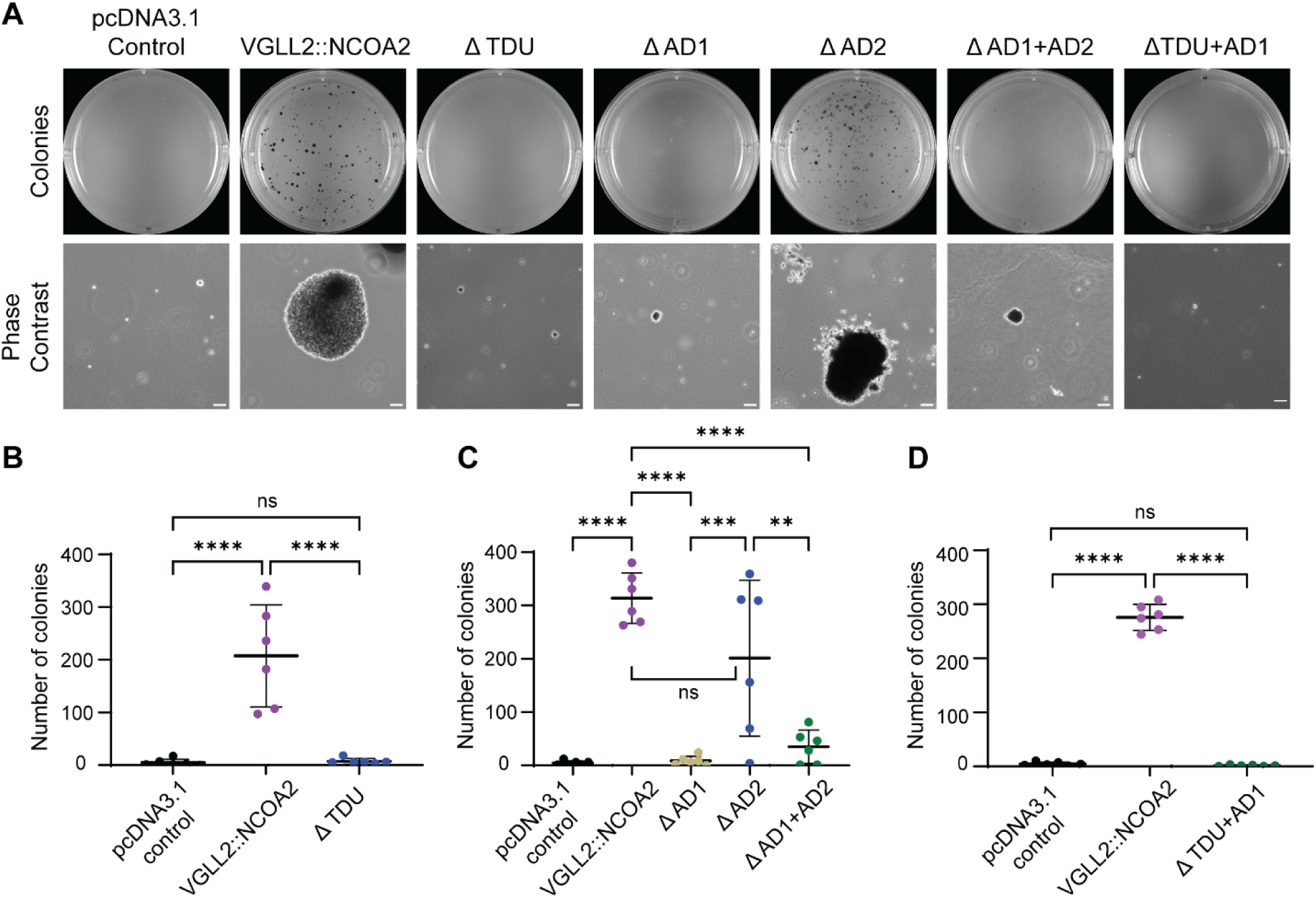
VGLL2::NCOA2 TDU and AD1 domains support oncogenic transformation of mouse myoblast cells. C2C12 mouse myoblast cells were transfected with pcDNA3.1 empty vector or pcDNA3.1 expressing 3’ end 3x FLAG-tag VGLL2::NCOA2 and domain deletion mutants. After two weeks of selection, they were plated for colony formation experiments in soft agar. (A) Representative images of the soft agar colony formation assay at two weeks post plating, including an image of the entire well and a phase contrast image. Scale bars: 100 µm. (B,C,D) Quantification of the number of colonies per well in a six-well plate for different transfected conditions. Data bars indicate mean ± SD. This was repeated three times independently with six technical replicates. The p values were calculated using a one-way ANOVA followed by Tukey’s multiple comparisons post hoc test. **p<0.01, ***p<0.001, ****p<0.0001, ns: not significant.

### VGLL2::NCOA2 TDU and AD1 domains alter transcriptomic profiles reflecting a less differentiated cell state

We used bulk RNA-seq to understand how the VGLL2::NCOA2 fusion-oncogene domains influence the transcriptomic landscape. We included the following groups transfected into C2C12 mouse myoblast cells grown in 2D: pcDNA control, full-length VGLL2::NCOA2, and all domain deletion mutants presented in Figure 1A. First, we performed principal component analysis and found a clear separation along PC1 between the full-length VGLL2::NCOA2 and AD2 deletion mutant as compared to the pcDNA control, and other domain deletion constructs, TDU, AD1, AD1+AD2, and TDU+AD1. Cells with the dual TDU+AD1 domain deleted construct separated along PC2 as a discrete transcriptomic profile (Figure 4A). This was also evident via hierarchical clustering (Figure 4B). These results indicate that, 1) the AD2 domain is dispensable for VGLL2::NCOA2 function in this context, and 2) the TDU and AD1 domains are key regulatory components of the VGLL2::NCOA2 transcriptome. Without the AD1 or TDU domains, the mutant VGLL2::NCOA2 expressing myoblasts cluster with a non-transformed pcDNA control mouse myoblast. Clustering is not solely dictated by VGLL2::NCOA2 mRNA levels, as the full-length VGLL2::NCOA2 and AD1 deletion mutant have the same mRNA levels of the fusion-oncogene expressed yet exhibit divergent colony formation capacity and transcriptomic signatures (Figure 3A,C and S5A). The full-length VGLL2::NCOA2 expressing cells compared to the pcDNA control cells up and down-regulate 4,171 and 4,062 genes, respectively, indicating the fusion-oncogene is a potent transcriptional regulator without bias for gene activation or repression (Figure S5B). The removal of any domain, or combination of domains, reduced the number of up or down-regulated genes (Figure S5B). When directly comparing all the mutant fusion-oncogenes with the full-length VGLL2::NCOA2 group, they ranged from ∼3700-4700 differentially expressed genes, except for the AD2 domain deleted group, which had a more muted impact on gene expression (Figure S5C).

**Figure 4:**
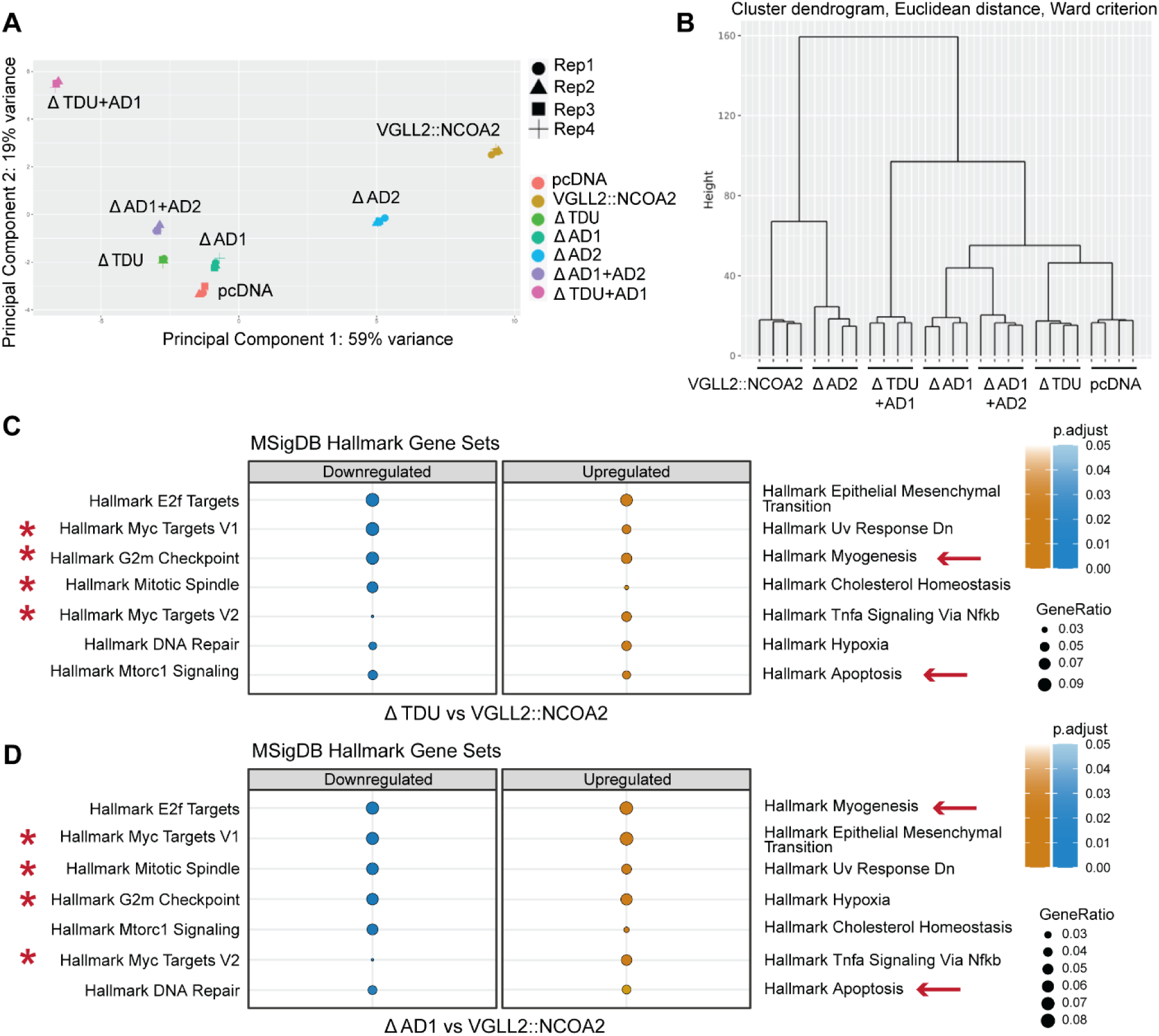
VGLL2::NCOA2 TDU and AD1 domains suppress skeletal muscle myogenesis and activate cell cycle gene sets. (A) Principal Component Analysis (after batch adjustment) of bulk RNA-seq expression data from C2C12 mouse myoblast controls or myoblasts expressing full-length VGLL2::NCOA2 or domain deletion mutants. (B) Hierarchical clustering of C2C12 transfection conditions based on normalized bulk RNA-seq expression data. (C, D) Hallmark gene sets enrichment plots detailing downregulated and upregulated gene sets in TDU domain deletion and AD1 domain deletion groups compared to the full-length VGLL2::NCOA2 condition. Reads were aligned to a custom mouse genome (mm39) that also included the human VGLL2::NCOA2 fusion-oncogene. Key downregulated gene sets are indicated by asterisks and upregulated gene sets by arrows.

We first confirmed our previous findings that VGLL2::NCOA2 driven zebrafish and mouse allograft tumors suppress myogenic pathways. We found this was also true in 2D systems, with a significant downregulation of hallmark myogenesis and apoptosis gene sets in VGLL2::NCOA2 expressing myoblasts compared to the pcDNA control myoblasts, whereas cell-cycle gene sets (Hallmark Myc targets v1/v2, mitotic spindle, G2M checkpoint) are upregulated (Figure S6A). This is likely attributed to VGLL2::NCOA2’s known role of suppressing skeletal muscle differentiation (27), and potentially priming processes important in transformation including cell proliferation and known oncogenic pathways. We then compared differentially regulated gene sets in all the domain deletion mutants of VGLL2::NCOA2 to the pcDNA control cells (Figure S6B-F). The hallmark myogenesis gene set was significantly downregulated in all the fusion-oncogene domain deletion mutants as compared to the pcDNA control group, indicating the mutants still have a capacity to suppress the myogenic process. Cell cycle gene sets were upregulated in the AD1, AD2, AD1+AD2 and TDU+AD1 domain deletions compared to the pcDNA control cells, while cell cycle gene sets were downregulated in the TDU domain deletion cells (Figure S6B-F). Thus, bulk RNA-seq analyses on 2D cells suggest that the VGLL2::NCOA2 TDU and AD1 domains are critical to regulate myogenesis and proliferative processes.

We then compared the control and deletion mutant forms of the fusion-oncogene to the full-length VGLL2::NCOA2 to understand how each domain contributed to discrete gene sets (Figure S7, 4C,D). We found that compared to the full-length VGLL2::NCOA2 expressing cells, the TDU and AD1 single domain deletion mutants had a significant upregulation of hallmark myogenesis and apoptosis gene sets and downregulation of cell-cycle gene sets (Figure 4C,D). These results suggest that VGLL2::NCOA2 utilizes the TDU and AD1 domains to orchestrate a more undifferentiated cell-state shift that includes the suppression of skeletal muscle differentiation and priming of proliferative processes. The AD2 domain did not have as big of an impact on the transcriptome. Deletion of the AD2 domain leads to an upregulation of the hallmark myogenesis and Myc targets v1/v2 cell-cycle gene sets compared to the full-length VGLL2::NCOA2 fusion-oncogene group. However, the magnitude of the enrichment for myogenesis gene sets is not as pronounced as with the TDU or AD1 deletion mutants (Figure S7B). Comparing the dual AD1+AD2 domain deletion mutant to the full-length VGLL2::NCOA2 fusion-oncogene group revealed that some gene sets now have reversed enrichment compared to the AD2 deletion (i.e. cell cycle gene sets and hallmark apoptosis) (Figure S7C). These changes reflect the pattern seen with the single AD1 deletion mutant, indicating that the AD1 domain is a more prevalent VGLL2::NCOA2 transcriptomic driver than the AD2 domain. The dual TDU+AD1 domain deletion has the most unique transcriptomic signature of all fusion-oncogene mutant groups (Figure 4A, Figure S7D). These findings suggest that in 2D cell culture, the TDU and AD1 domains could work cooperatively in an independent mechanism from the individual domain deletions to support a distinct cell state.

### VGLL2::NCOA2’s TDU and AD1 domains support tumorigenesis in a mouse allograft model

Previously, we showed that untagged VGLL2::NCOA2 expressed in mouse myoblasts readily generated fast-growing tumors in a mouse allograft model. In the same study, control C2C12 mouse myoblasts transfected with a pcDNA 3.1 backbone resulted in small tumor lesions after orthotopic injection into skeletal muscle (27). Here, we used a similar approach and allografted C2C12 mouse myoblasts stably expressing either full-length VGLL2::NCOA2 or TDU, AD1, and TDU+AD1 domain deletions intramuscularly into severe combined immunodeficient (Icr-SCID) mice. Mice allografted with mouse myoblasts containing the full-length VGLL2::NCOA2 FLAG tagged fusion-oncogene rapidly formed tumors with an average initial onset of 21 days post injection, consistent with our previously published work using an untagged VGLL2::NCOA2 (27). Mice allografted with mouse myoblasts containing the VGLL2::NCOA2 fusion-oncogene with TDU or AD1 domain deletions had a significant delay in tumor onset with initial detection occurring ∼63 days post injection. Mice allografted with mouse myoblasts containing the VGLL2::NCOA2 fusion-oncogene with dual TDU+AD1 domain deletions had a further delay in tumor onset occurring ∼73 days post injection (Left panel, Figure 5A). Plotting the cumulative tumor volume as a function of days post injection revealed that full-length VGLL2::NCOA2 tumors were more aggressive and reached tumor endpoint (tumor size of 1000 mm^3^) at a faster rate compared to the tumors formed by the VGLL2::NCOA2 TDU, AD1 and TDU+AD1 domain deletion constructs (Right panel, Figure 5A). The full-length VGLL2::NCOA2 tumors reached endpoint ∼39 days, whereas at the experimental endpoint of 122 days, the tumor volume in the TDU deletion mutant was more than 500 mm^3^ for 6 out of the 8 mice, and in the AD1 deletion mutant the tumor volume was greater than 500 mm^3^ in all 7 mice. In the TDU+AD1 deletion mutant, 5 out of 7 mice had tumors with volumes more than 500 mm^3^ (Figure 5B,C). The allograft experiment was repeated, and we observed similar trends in tumor volume compared to the first trial (Figure S8A-C). These results indicate that the TDU and AD1 domains support optimal VGLL2::NCOA2 tumorigenesis and that removing these domains, and likely their protein interactions, suppresses fusion-driven tumor onset and growth.

**Figure 5:**
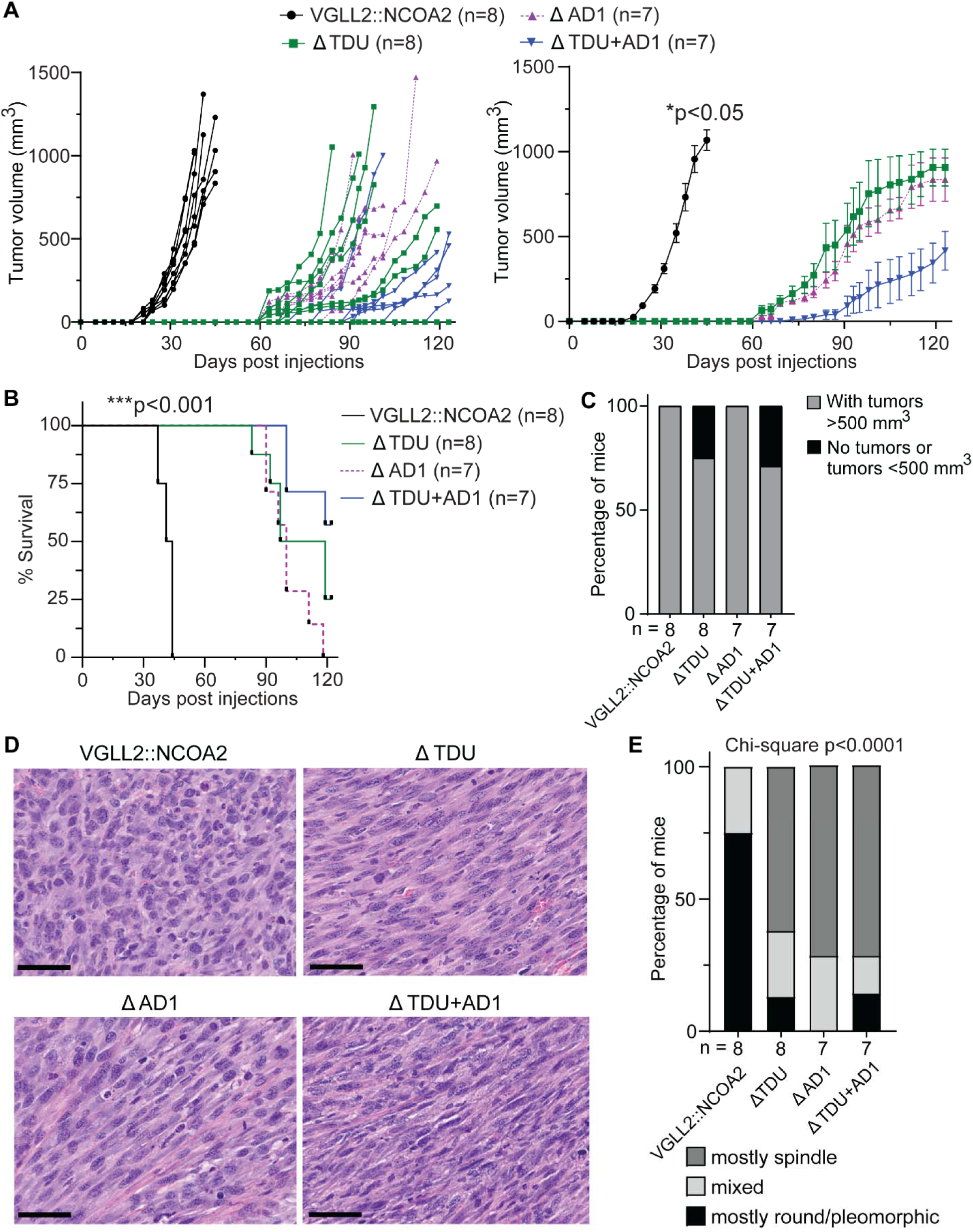
VGLL2::NCOA2 TDU and AD1 domains support fusion-driven tumor formation in a mouse allograft model. (A) Tumor leg volume is plotted for each individual mouse as a function of days post injection (left). Cumulative mean tumor volume of all the tumor-bearing mice as a function of days post injection (right). Data indicate mean ± standard error of mean (SEM). p values were calculated by comparing the mean days of tumor onset using a Kruskal Wallis non-parametric ANOVA test corrected for multiple comparisons using the 2-stage linear step-up procedure of Benjamini, Krieger, and Yekutieli. *p<0.05 with respect to comparing the full-length VGLL2::NCOA2 condition to all domain deletion mutant allograft tumors. (B) Survival plot indicating percentage of mice reaching tumor or humane endpoint for each condition as a function of days post injections. Cumulative statistical analysis was performed using Log-rank (Mantel-Cox) test and individual tests between two groups were performed using Log-rank (Mantel-Cox) test and confirmed using Gehan-Breslow-Wilcoxon test. ***p<0.001 with respect to comparing the full-length VGLL2::NCOA2 condition to all domain deletion mutant allograft tumors. (C) Stacked bar plot indicating the number of mice with tumors greater than 500 mm^3^ volume, and number of mice with no tumors and tumors smaller than 500 mm^3^ volume. (D) Representative hematoxylin-eosin (H&E) staining from a C2C12 VGLL2::NCOA2 full-length allograft and a C2C12 VGLL2::NCOA2 with domain deletions of TDU, AD1 and TDU+AD1 tumor allografts. Scale bars: 50 μm. (E) Pathological assessment of tumor cell morphology based on hematoxylin and eosin (HE)- stained slides in a semi-quantitative manner, using a three-tiered system: mostly (>75%) round/pleomorphic cells, mostly (>75%) spindle cells, or a combination of the above (mixed). Statistical significance was determined using a Chi-square test.

### VGLL2::NCOA2 TDU and AD1 domains support more aggressive histological features in fusion-driven infantile rhabdomyosarcoma

We quantified histopathological differences in the full-length VGLL2::NCOA2 fusion-oncogene and domain deletion tumors using hematoxylin and eosin (H&E) staining and evaluation by a pediatric sarcoma expert pathologist (J.C.; Figure 5D). There was a higher proportion of cells exhibiting round or pleomorphic morphology in the full-length VGLL2::NCOA2 tumors compared to domain deletion tumors (Figure 5D,E, S8D). This correlates well with our published untagged VGLL2::NCOA2 allograft model. In this system, full-length VGLL2::NCOA2 tumors exhibited more pleomorphic morphology compared to sections taken from control allografted C2C12-pcDNA lesions, which displayed primarily a spindle cell morphology (27). In the VGLL2::NCOA2 domain deletion tumors, the histology was predominantly a spindle cell morphology (Figure 5D,E, S8D). Additionally, the full-length VGLL2::NCOA2 tumors had a significantly larger area and higher percentage of necrosis compared to the tumors in the domain deletion groups, providing another indication that the full-length tumors were more aggressive (Figure 6A,B, S8E,F). These trends were consistent across the independent allograft trials. These same histological features of pleomorphic cell morphology and tumor necrosis were also observed in patients with VGLL2::NCOA2 tumors that had local progression and tumor relapse (7).

**Figure 6:**
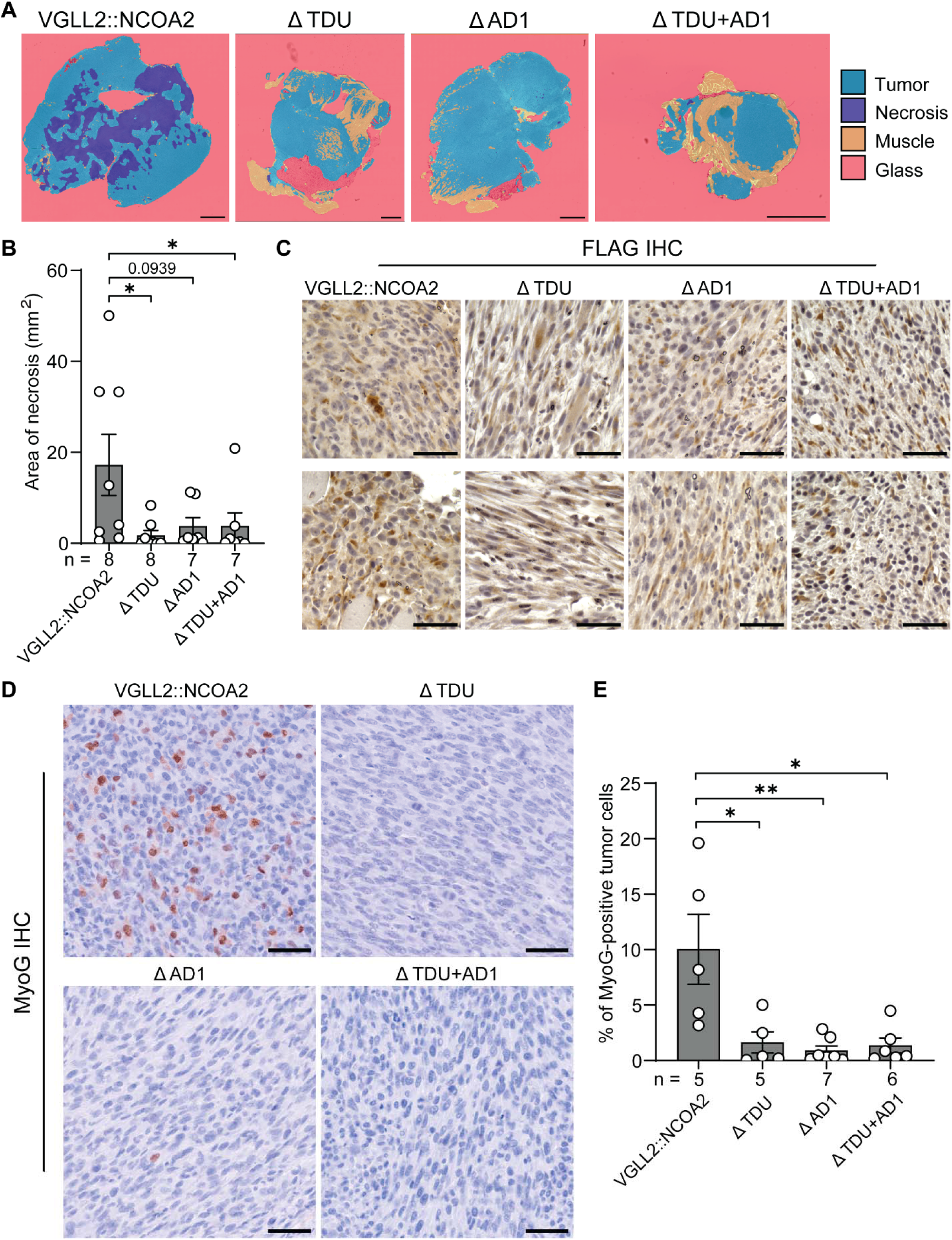
VGLL2::NCOA2 TDU and AD1 domains support more aggressive histological features in fusion-driven mouse allograft tumors. (A) Heatmaps generated by the HALO platform indicating regions of tumor, necrosis and normal skeletal muscle on the allograft sections for different fusion-oncogene or mutant conditions. Scale bars: 2 mm (B) Quantification of necrosis areas on hematoxylin and eosin (HE)-stained slides was performed using the HALO image analysis platform and the HALO AI module (version 3.5.3577) from Indica Labs (Albuquerque, NM, USA). Data indicate mean ± SEM. Statistical analysis comparing mean area of necrosis in the full-length VGLL2::NCOA2 group to the domain deletion mutants was performed using the Mann Whitney non-parametric test. *p<0.05. (C, D) Representative images of FLAG and MyoG tumor immunohistochemistry in the VGLL2::NCOA2, TDU deletion, AD1 deletion, and TDU and AD1 deletion conditions. Scale bars: 50 μm. (E) Quantification of the percentage of myogenin-positive tumor nuclei on stained slides was performed using the HALO image analysis platform and the HALO AI module (version 3.5.3577) from Indica Labs (Albuquerque, NM, USA). Data indicate mean ± SEM. Statistical analysis comparing mean %myogenin-positive tumor cells in the full-length VGLL2::NCOA2 group to the domain deletion mutants was performed using the Mann Whitney non-parametric test. *p<0.05, **p<0.01.

Next, we evaluated the protein expression of FLAG as a proxy for VGLL2::NCOA2 tumor cells, myogenin (MyoG), which is a RMS diagnostic marker, and all myosin heavy chains (MYH), which is indicative of terminal muscle differentiation. We performed FLAG immunohistochemistry and saw that all of the tumors expressed variable FLAG staining, indicating that the VGLL2::NCOA2 fusion-oncogene or domain deletion mutants are expressed (Figure 6C). We also performed quantitative immunohistochemistry of MyoG, which in C2C12 cells is absent in cycling myoblasts, increases in expression when myoblasts are crowded and withdraw from the cell cycle to initiate differentiation, and then decreases in expression as cells undergo terminal differentiation into multi-nucleated myotubes (48). Previously, we showed that C2C12 pcDNA control myoblasts allografted at the injection site heterogeneously express MyoG likely due to the crowding of the cells and a subset withdrawing from the cell cycle. We found that the full-length VGLL2::NCOA2-driven tumors heterogeneously express MyoG in ∼10% of the cells. However, tumors in the domain deletion groups had a significant decrease in MyoG protein expression (Figure 6D,E). This pattern was trending in the replicate, and when the data across trials are combined, the full-length VGLL2::NCOA2 condition is statistically significant from the domain deletion mutants (Figure S9A,B). Myosin heavy chain is a terminal differentiation marker and structural protein and is typically not detected in rhabdomyosarcoma. We performed myosin heavy chain immunohistochemistry on tumor sections and found no detectable protein in full-length VGLL2::NCOA2 tumors, and some elongated fibers or detectable myosin heavy chain expression in the domain deletion mutant tumors (Figure S9C). However, this could represent entrapped normal skeletal muscle. These observations support that the tumors in the VGLL2::NCOA2 domain deletion groups are histologically different and represent a less aggressive and more differentiated disease.

## Discussion

The primary genetic drivers in fusion-driven rhabdomyosarcoma involve gene fusions of transcription factors and co-activators that are critical to skeletal muscle development; including, PAX3::FOXO1 and PAX7::FOXO1 in alveolar rhabdomyosarcoma (49–52), and VGLL2::NCOA2, SRF::NCOA2, VGLL2::CITED2, and TEAD1::NCOA2 in the recently described infantile spindle cell/sclerosing rhabdomyosarcoma (5–10). Under normal developmental conditions, these 5’ and 3’ fusion partners orchestrate transcription factor networks that drive skeletal muscle specification, migration, and differentiation (16–22, 53–58). In our published work, we showed that the VGLL2::NCOA2 fusion is indeed an oncogene that generates aggressive tumors in both a mouse myoblast allograft model, and a transgenic zebrafish model. These models have high fidelity to the human disease at the histological and transcriptomic levels (27). In this current study, we build on these findings and detail the VGLL2::NCOA2 fusion-oncogene and the functional contributions of its protein domains to tumorigenesis using a structure-function approach. We find that the TDU domain of VGLL2 and the AD1 domain of NCOA2 direct fusion-driven transcriptomic signatures, suppressing myogenic differentiation, supporting transformation capacity, and promoting more aggressive disease. This is true in both cell-based and mouse allograft systems. Notably, the 5’ or 3’ gene fusion partners all have functional redundancy or overlap (16–19, 53–55), suggesting this could also be generally true in fusion-driven infantile RMS and that these findings could be useful for understanding core disease dependencies.

We show that the TDU and AD1 domains are driver components in mediating VGLL2::NCOA2 transformation and suppressing skeletal muscle differentiation. Individually, the removal of these domains delays tumor formation in our mouse allograft model, and the dual TDU+AD1 domain removal further suppresses tumor formation. However, in all allograft contexts, tumors do eventually form, indicating there are additional interactions and mechanisms that support tumorigenesis outside of the domains assessed here. The C terminus of NCOA2 is intrinsically disordered (33, 59) and forms molecular condensates in cancer and normal contexts (22, 60–62). The VGLL family can contain intrinsically disordered regions (63, 64), but the evidence for condensate formation is not as compelling as for NCOA2. The mutations we introduced to VGLL2::NCOA2 still retained portions of NCOA2 outside of AD1 and AD2. Ectopic expression of HEY1::NCOA2 and PAX5::NCOA5 in HeLa cells forms molecular condensates that induce aberrant gene transcription or inappropriately sequester regulators away from their normal function (62). NCOA2 also directly binds acetyltransferase p300, p300-associated factor PCAF, and transcription factor MEF2 to form molecular condensate-like nuclear puncta in cultured cells and regulate skeletal muscle differentiation (22, 60, 61). In C2C12 mouse myoblasts, p300 interacts with transcription factor MyoD, which leads to cell-cycle arrest, one of the earlier steps in the differentiation process (65). This direct binding of VGLL2::NCOA2 and endogenous p300 is retained after ectopic fusion-oncogene expression in human HEK293T cells (28). Thus, it is possible that the VGLL2::NCOA2 fusion-oncogene utilizes either the TDU domain, AD1 domain, or both, to localize into the nucleus resulting in molecular condensate formation that likely includes TEAD1, p300, and other interacting partners. Further studies are required to confirm global interacting nuclear partners of VGLL2::NCOA2 to provide a more focused epigenetic perspective on transcriptional regulation.

We also find evidence that NCOA2 domain dependencies are in-part conserved across other NCOA2-fused tumors. NCOA2 is a common 3’ fusion partner within and outside of RMS (66), including other cancers such as HEY1::NCOA2 in mesenchymal chondrosarcoma and MOZ::NCOA2 in acute myeloid leukemia (67, 68). We compared our results to a structure-function study of the PAX3::NCOA2 fusion-oncogene (69), a genetic driver found in alveolar rhabdomyosarcoma and biphenotypic sinonasal sarcoma (70). In NIH3T3 mouse fibroblasts, PAX3::NCOA2 mutants with deletion of the AD1 domain had a significant reduction in colony formation compared to cells with the full-length PAX3::NCOA2. This is similar to our findings. However, the study also found that a PAX3::NCOA2 AD2 domain deletion mutant decreased colony formation, but to a lesser extent than the AD1 domain-deletion group (69). In our data, bulk RNA-seq transcriptomic signatures suggest that in mouse myoblast cells, the VGLL2::NCOA2 AD2 domain deletion mutant is similar to the full-length VGLL2::NCOA2 group. Further, C2C12 mouse myoblasts expressing the VGLL2::NCOA2 AD2 deletion mutant form the same number of colonies as the full-length VGLL2::NCOA2 group. This suggests that the VGLL2::NCOA2 AD2 domain is not a driver of its tumorigenic activity. Key differences in our studies are the fusion-oncogene 5’ partner, cellular context used for the assessment of transformation capacity, and the definition of the AD2 domain and the amino acids it spans. We removed 177 amino acids, consistent with the literature for the VGLL2::NCOA2 AD2 deletion mutant (32, 33), compared to the approximately ∼33 amino acids for the PAX3::NCOA2 AD2 deletion mutant. Given that the NCOA2 AD2 domain is known to interact with histone methyltransferase CARM1/PRMT1 (26), there could be a role that is dispensable for tumor initiation, but is important in tumor maintenance and evolution (71, 72). Our findings support that AD1 is a key contributor to NCOA2-fused fusion-oncogene activity, and we hypothesize that targeting the AD1 domain and its interacting partners represents a shared dependency in NCOA2 fusion-driven cancers.

Previous studies have investigated targeting TEAD1 and p300/CBP, two known TDU and AD1 domain interacting proteins, in other cancers. This represents a potential therapeutic approach in infantile spindle cell/sclerosing RMS and could be required given that single domain deletions did not completely abrogate tumor formation in our mouse allograft model. p300 has been proposed as a cooperating gene and therapeutic target in a broad range of cancers, including pediatric or adolescent young adult (AYA) fusion-driven sarcomas such as PAX3::FOXO1 RMS (73), SS18::SSX synovial sarcoma (bioRxiv 2026.01.26.701706, bioRxiv 2026.01.26.701739), CIC::DUX4 sarcoma, and EWS::FLI1 Ewing sarcoma (74, 75). In CIC::DUX4 fusion-driven sarcoma, targeting p300/CBP using a small molecule inhibitor, iP300w, downregulated CIC::DUX4 target genes and suppressed cell proliferation in patient cells and xenograft models (74). Similar effects were seen with iP300w treatment in EWS::FLI1-mediated Ewing sarcoma (75). A less potent small molecule p300 inhibitor, A485, suppressed colony formation and tumor growth in mouse allograft models of VGLL2::NCOA2 and TEAD1::NCOA2-driven infantile spindle cell/sclerosing RMS (28). TEAD1 and co-option of TEAD transcriptional programs could also represent a therapeutic avenue and have shown proof of concept efficacy in PAX3::FOXO1-driven rhabdomyosarcoma. Treatment of a PAX3::FOXO1 RMS patient cell line with verteporfin and protoporphyrin IX, inhibitors of the Hippo signaling pathway, suppressed TEAD transcriptional activity, and decreased cell and tumor growth in culture and in mouse xenograft models (76). Future studies will address the direct binding of TEAD1 and p300/CBP with VGLL2::NCOA2 structural domains and specific residues. Further, we will link how VGLL2::NCOA2 and domain mutants modify the genome-wide changes in DNA binding of p300/CBP and TEAD1 to understand genetic cooperation and direct and indirect transcriptomic consequences. Overall, elucidating the transcriptional and epigenetic mechanisms in VGLL2::NCOA2 driven RMS will identify direct druggable targets that both promote skeletal muscle differentiation and suppress oncogenic activity.

## Summary

Infantile spindle cell/sclerosing RMS is driven by gene fusions, the most common of which is VGLL2::NCOA2. In this work, we use a structure function-based approach to delineate the roles of VGLL2::NCOA2’s protein domains to engage skeletal muscle programs and support tumorigenesis. The VGLL2 TDU, and NCOA2 AD1 and AD2 domains, normally interact with transcription factors and epigenetic regulators such as TEAD1, p300/CBP and CARM1/PRMT1; however, their regulatory roles in VGLL2::NCOA2 oncogenic activity are not fully understood. Using a combination of cell culture and mouse allograft studies, we find that the TDU and AD1 domains are critical contributors to initiating transformation, supporting tumor growth, and suppressing skeletal muscle terminal differentiation. Targeting these domains, particularly a dual approach of targeting TDU and AD1 protein interactions, will facilitate identification of novel strategies to inhibit VGLL2::NCOA2 activity and potentially be used as therapeutic opportunities. More broadly, we anticipate these targets and strategies could have applicability to the spectrum of NCOA2-based fusion-oncogenes identified in infantile RMS and other cancer types.

## Acknowledgements

We thank the Nationwide Children’s Hospital (NCH) Animal Resources Core, the NCH Tumor Core, the Steve and Cindy Rasmussen Institute for Genomic Medicine, Genomics Services Laboratory, NCH Histopathology, Processing, and Banking Core, and the NCH High-Performance Computing Group for assistance in maintaining and using the NCH cluster. We are grateful to all members of the Kendall lab for helpful discussions.

## Author Contributions

**C. S. Sankhe**: Conceptualization, Data curation, Formal analysis, Investigation, Methodology, Validation, Visualization, Writing - original draft, Writing – review & editing. **D. Calderon**: Conceptualization, Data curation, Formal analysis, Investigation, Methodology, Writing - original draft, Writing – review & editing. **C. Taslim**: Data curation, Investigation, Resources, Software, Validation, Writing – review & editing. **M. Sammons, L. Ryan, A. C. Gross**: Investigation, Methodology, Validation, Writing – review & editing. **K. M. Silvius**: Methodology, Writing – review & editing. **O. Delattre**: Resources, Writing – review & editing. **J. Cyrta**: Funding acquisition, Investigation, Methodology, Validation, Visualization, Writing – review & editing**. R. Roberts**: Methodology, Resources, Supervision, Writing – review & editing. **G. C. Kendall**: Conceptualization, Data curation, Formal analysis, Funding acquisition, Investigation, Methodology, Project administration, Resources, Software, Supervision, Writing - original draft, Writing – review & editing

