## Supplementary Figures for "Structure-Function Analysis of an Infantile Rhabdomyosarcoma Fusion-Oncogene Reveals Domain Requirements for Tumorigenesis"

### Title

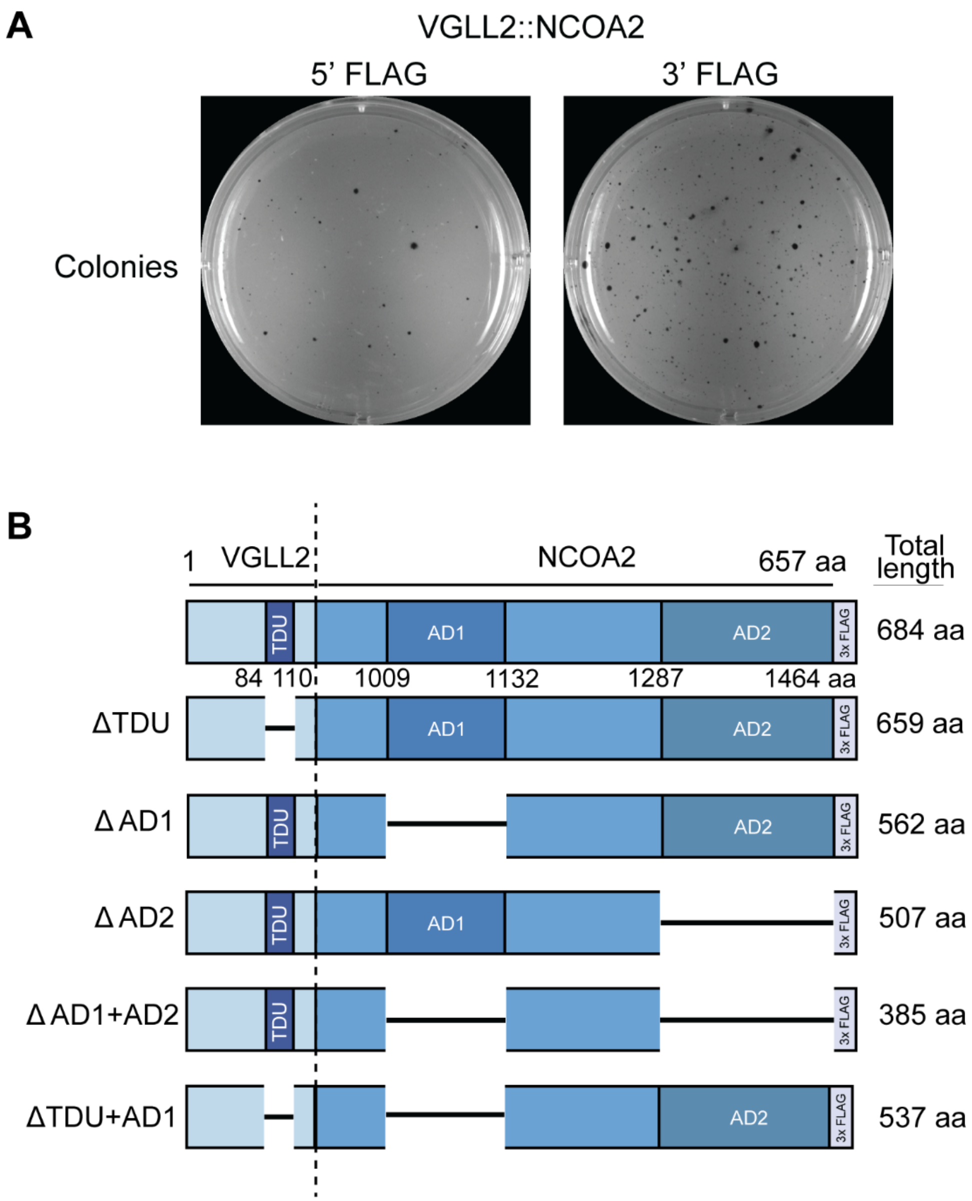


**Supplementary Figure S1: Determining the optimal FLAG tag location and specific domain deletion mutants used in experiments.**

(A) Representative images of wells containing colonies in soft agar taken after 14 days post plating for C2C12 cells transfected with the VGLL2::NCOA2 containing a 3x FLAG tag on either the 5' end or 3' end.

(B) Schematic depicting full-length and mutant VGLL2::NCOA2 fusion-oncogene constructs with deleted TDU, AD1, AD2, AD1+AD2 domains, and TDU+AD1 domains. The numbers indicate the amino acid position where the deletion was performed for the respective domains. The exact mutations are as follows: the TDU domain mutant deletion is from amino acids 85 to 109, the AD1 domain mutant deletion is from amino acids 1010 to 1131, and the AD2 mutant deletion is from amino acids 1288 to 1464.

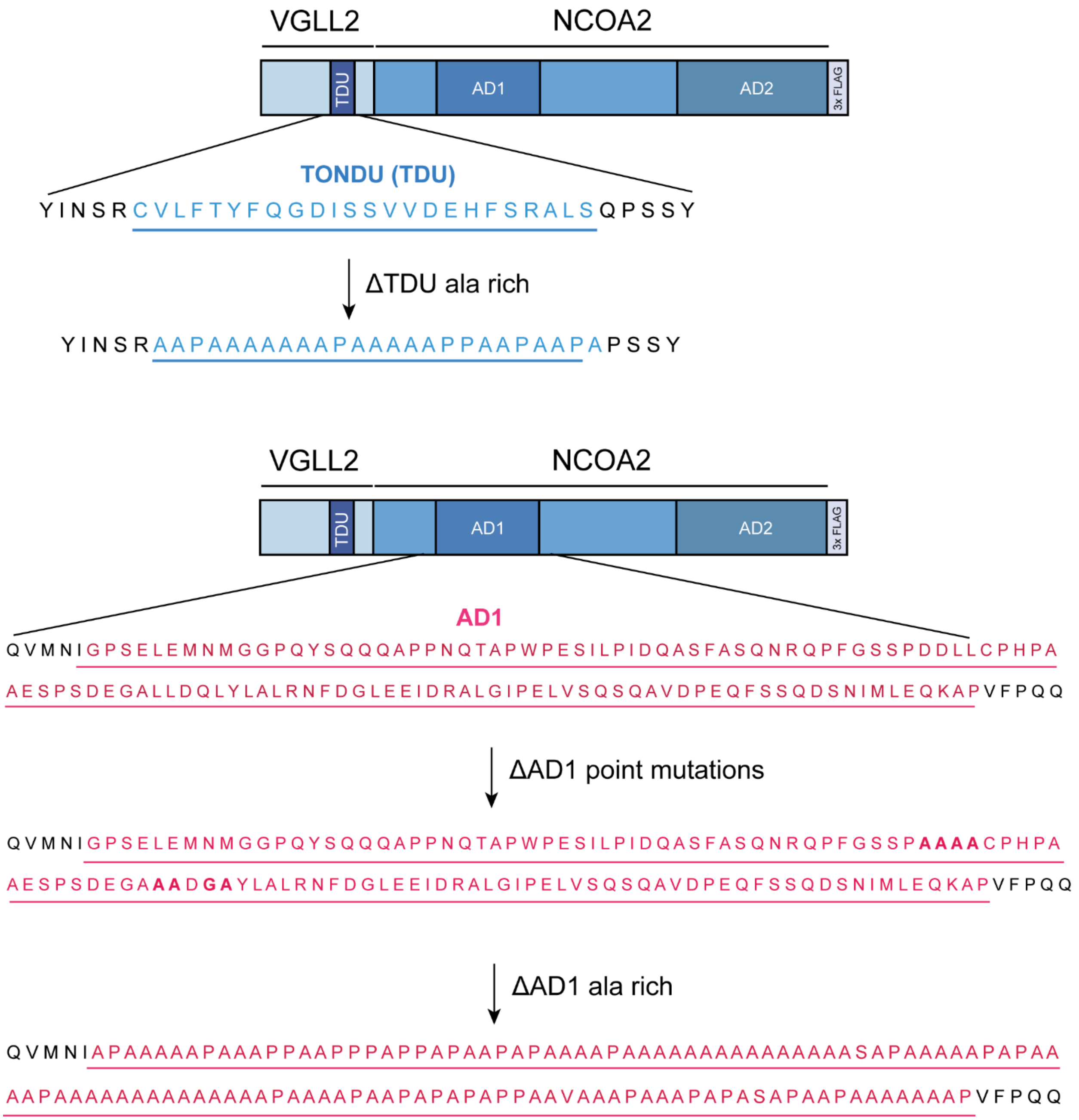


**Supplementary Figure S2: Schematic depicting complementary mutations in the TDU and AD1 domains including an alanine stretch substitution in the TDU and AD1 domains and point mutations in the AD1 domain.**

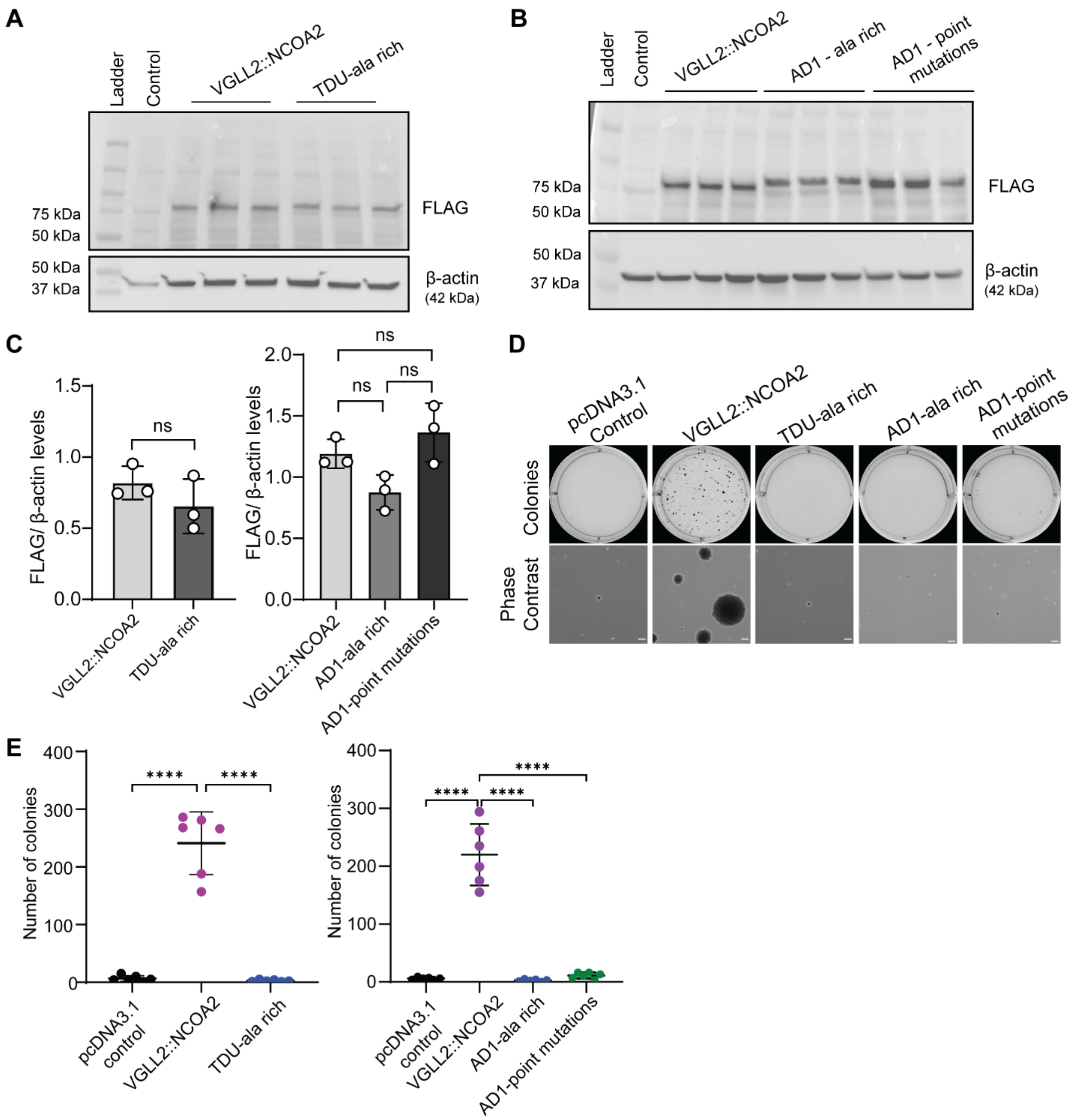


**Supplementary Figure S3: VGLL2::NCOA2 TDU and AD1 domain alanine-rich or point mutations result in comparable protein expression to the full-length VGLL2::NCOA2 fusion, yet do not generate colonies in colony formation assays.**

(A, B) Western blot of all three technical replicates of whole-cell protein lysates extracted from C2C12 cells transfected with full-length VGLL2::NCOA2, point mutated TDU domain substituted with the alanine(Ala)-rich sequence, point mutated AD1 domain substituted with the alanine(Ala)-rich sequence or mutated at locations in the domain sequence, and probed for FLAG and loading control β-actin. Control sample was the whole-cell lysate extracted from the cells transfected with the pcDNA3.1 vector.

(C) Quantification of the relative levels of FLAG in C2C12 transfected cells via densitometry analysis from the western blots in Panel (A, B). Data indicate mean ± SD for n=3 technical replicates. The significance test used was T-test with Welch's correction and Kruskal-Wallis non-parametric ANOVA with Dunn's multiple comparisons post hoc test. ns: not significant.

(D) Representative images of wells containing colonies in soft agar taken after 14 days post plating for each transfection group along with representative phase contrast images for each condition. Scale bars: 100 μm.

(E) Quantification of the number of colonies per well in a 6-well plate for different transfected conditions. Data indicate mean ± SD. This was repeated two times independently with six technical replicates. The p values were calculated using a one-way ANOVA followed by Tukey's multiple comparisons post hoc test. ****$p<0.0001$.

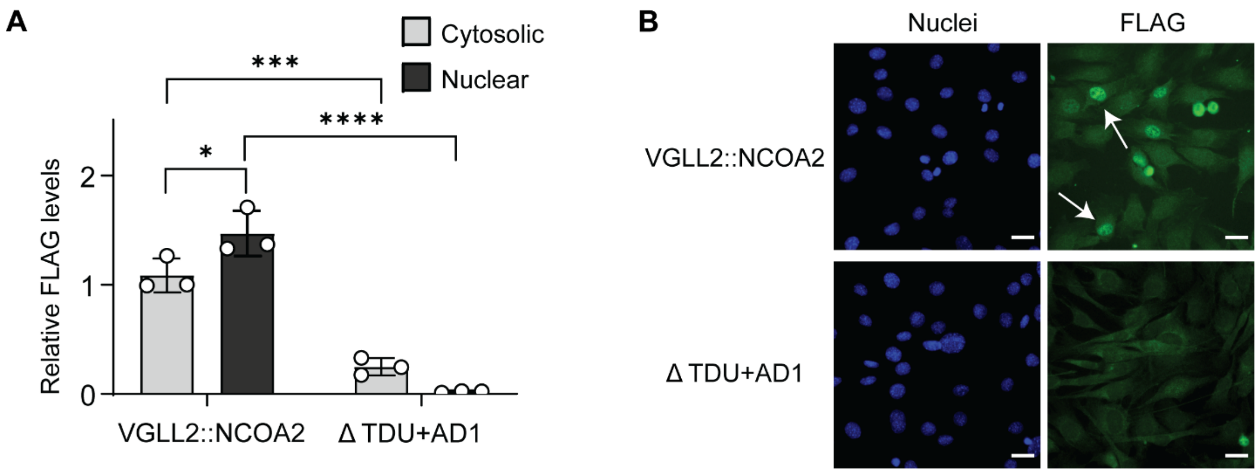


**Supplementary Figure S4: VGLL2::NCOA2 TDU+AD1 dual domain deletion results in decreased fusion-oncogene nuclear localization.**

(A) Quantification of relative nuclear and cytosolic FLAG levels in C2C12 transfected cells via densitometry analysis from the western blot in Panel (G) in Figure 1. Data indicate mean ± SD for n=3 replicates. The p values were calculated using a one-way ANOVA followed by Tukey's multiple comparisons post hoc test. *$p<0.05$, ***$p<0.001$, ****$p<0.0001$.

(B) Immunofluorescence staining of FLAG in transfected C2C12 cells with full-length fusion-oncogene and mutant TDU+AD1 domain deleted fusion-oncogene. White arrows indicate cells with nuclear localized FLAG (VGLL2::NCOA2). Scale bars: 25 μm.

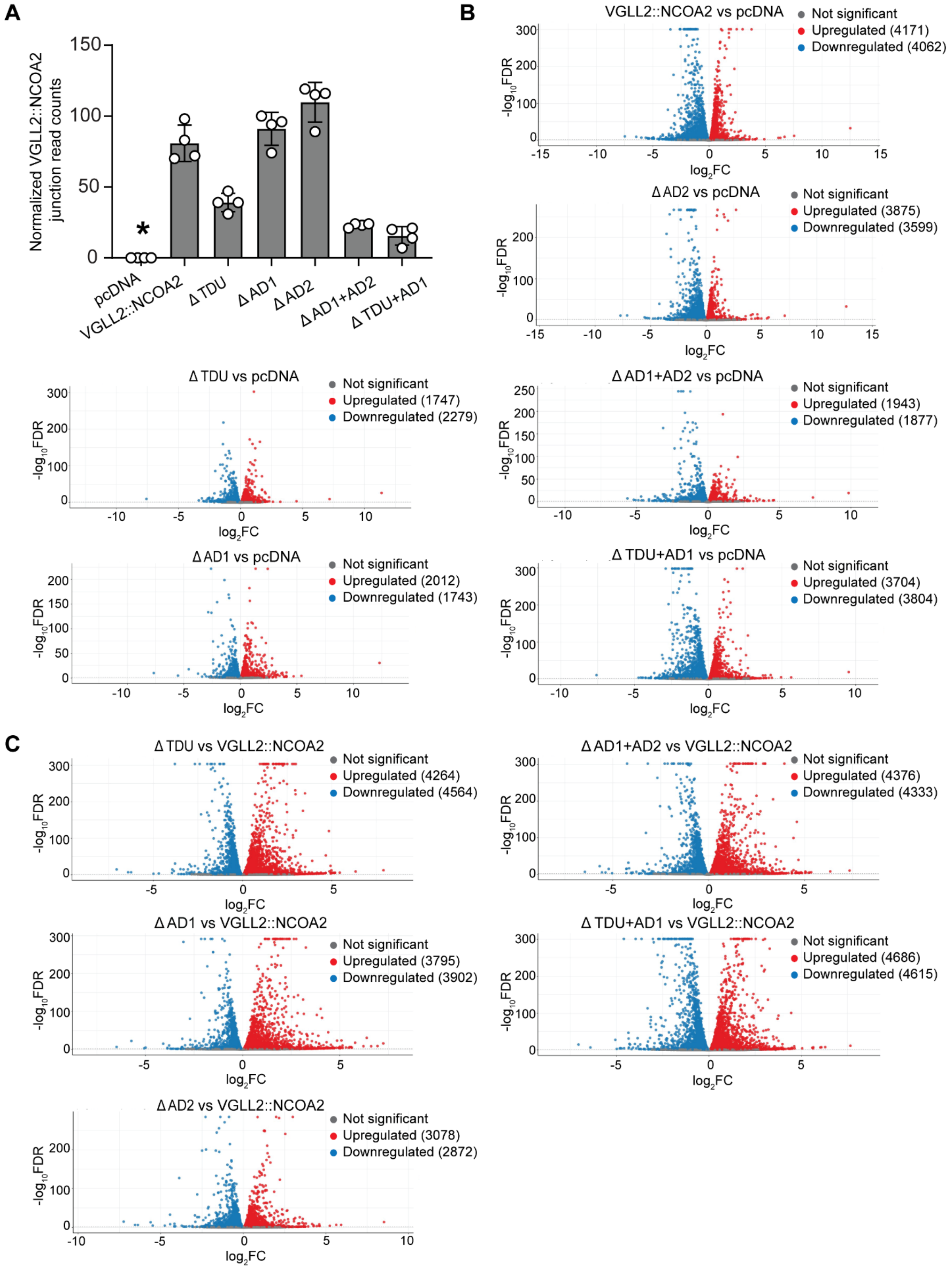

A
Normalized VGLL2::NCOA2 junction read counts
pcDNA
VGLL2::NCOA2
Δ TDU
Δ AD1
Δ AD2
Δ AD1+AD2
Δ TDU+AD1
B
VGLL2::NCOA2 vs pcDNA
Not significant
Upregulated (4171)
Downregulated (4062)
Δ AD2 vs pcDNA
Upregulated (3875)
Downregulated (3599)
Δ TDU vs pcDNA
Upregulated (1747)
Downregulated (2279)
Δ AD1+AD2 vs pcDNA
Upregulated (1943)
Downregulated (1877)
Δ AD1 vs pcDNA
Upregulated (2012)
Downregulated (1743)
Δ TDU+AD1 vs pcDNA
Upregulated (3704)
Downregulated (3804)
C
Δ TDU vs VGLL2::NCOA2
Upregulated (4264)
Downregulated (4564)
Δ AD1+AD2 vs VGLL2::NCOA2
Upregulated (4376)
Downregulated (4333)
Δ AD1 vs VGLL2::NCOA2
Upregulated (3795)
Downregulated (3902)
Δ TDU+AD1 vs VGLL2::NCOA2
Upregulated (4686)
Downregulated (4615)
Δ AD2 vs VGLL2::NCOA2
Upregulated (3078)
Downregulated (2872)
-log10FDR
log2FC

**Supplementary Figure S5: Volcano plots and VGLL2::NCOA2 fusion-oncogene and mutant levels from bulk RNA-seq analysis.**

(A) Normalized VGLL2::NCOA2 junction read counts for different transfection groups. * indicates normalized junction sequence read counts are significantly upregulated in the full-length VGLL2::NCOA2 condition and all domain deletion mutants compared to the pcDNA control group. Reads were aligned to a custom mouse genome (mm39) that also included the human VGLL2::NCOA2 fusion-oncogene junction sequence.

(B,C) Volcano plots listing the number of differentially upregulated and downregulated genes comparing all the transfection groups to pcDNA control transfection condition (B), and comparing all the domain-deleted VGLL2::NCOA2 fusion-oncogene groups to the full-length VGLL2::NCOA2 condition (C). Reads were aligned to a custom mouse genome (mm39) that also included the human VGLL2::NCOA2 fusion-oncogene.

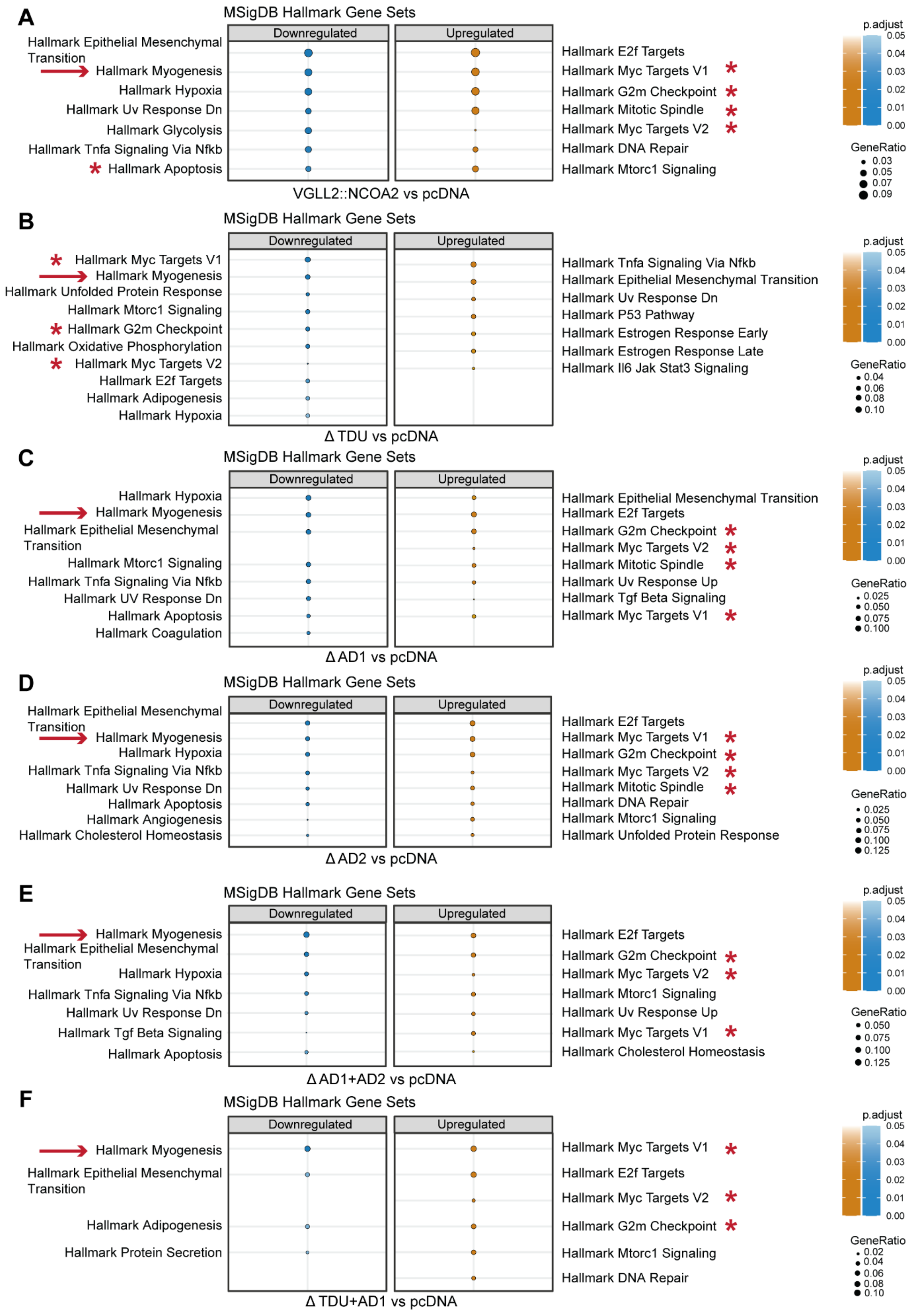
A
MSigDB Hallmark Gene Sets
Downregulated
Upregulated
Hallmark Epithelial Mesenchymal Transition
Hallmark Myogenesis
Hallmark Hypoxia
Hallmark Uv Response Dn
Hallmark Glycolysis
Hallmark Tnfa Signaling Via Nfkb
Hallmark Apoptosis
Hallmark E2f Targets
Hallmark Myc Targets V1
Hallmark G2m Checkpoint
Hallmark Mitotic Spindle
Hallmark Myc Targets V2
Hallmark DNA Repair
Hallmark Mtorc1 Signaling
VGLL2::NCOA2 vs pcDNA
p.adjust
0.05
0.04
0.03
0.02
0.01
0.00
GeneRatio
0.03
0.05
0.07
0.09
B
MSigDB Hallmark Gene Sets
Downregulated
Upregulated
Hallmark Myc Targets V1
Hallmark Myogenesis
Hallmark Unfolded Protein Response
Hallmark Mtorc1 Signaling
Hallmark G2m Checkpoint
Hallmark Oxidative Phosphorylation
Hallmark Myc Targets V2
Hallmark E2f Targets
Hallmark Adipogenesis
Hallmark Hypoxia
Hallmark Tnfa Signaling Via Nfkb
Hallmark Epithelial Mesenchymal Transition
Hallmark Uv Response Dn
Hallmark P53 Pathway
Hallmark Estrogen Response Early
Hallmark Estrogen Response Late
Hallmark Il6 Jak Stat3 Signaling
Δ TDU vs pcDNA
p.adjust
0.05
0.04
0.03
0.02
0.01
0.00
GeneRatio
0.04
0.06
0.08
0.10
C
MSigDB Hallmark Gene Sets
Downregulated
Upregulated
Hallmark Hypoxia
Hallmark Myogenesis
Hallmark Epithelial Mesenchymal Transition
Hallmark Mtorc1 Signaling
Hallmark Tnfa Signaling Via Nfkb
Hallmark UV Response Dn
Hallmark Apoptosis
Hallmark Coagulation
Hallmark Epithelial Mesenchymal Transition
Hallmark E2f Targets
Hallmark G2m Checkpoint
Hallmark Myc Targets V2
Hallmark Mitotic Spindle
Hallmark Uv Response Up
Hallmark Tgf Beta Signaling
Hallmark Myc Targets V1
Δ AD1 vs pcDNA
p.adjust
0.05
0.04
0.03
0.02
0.01
0.00
GeneRatio
0.025
0.050
0.075
0.100
D
MSigDB Hallmark Gene Sets
Downregulated
Upregulated
Hallmark Epithelial Mesenchymal Transition
Hallmark Myogenesis
Hallmark Hypoxia
Hallmark Tnfa Signaling Via Nfkb
Hallmark Uv Response Dn
Hallmark Apoptosis
Hallmark Angiogenesis
Hallmark Cholesterol Homeostasis
Hallmark E2f Targets
Hallmark Myc Targets V1
Hallmark G2m Checkpoint
Hallmark Myc Targets V2
Hallmark Mitotic Spindle
Hallmark DNA Repair
Hallmark Mtorc1 Signaling
Hallmark Unfolded Protein Response
Δ AD2 vs pcDNA
p.adjust
0.05
0.04
0.03
0.02
0.01
0.00
GeneRatio
0.025
0.050
0.075
0.100
0.125
E
MSigDB Hallmark Gene Sets
Downregulated
Upregulated
Hallmark Myogenesis
Hallmark Epithelial Mesenchymal Transition
Hallmark Hypoxia
Hallmark Tnfa Signaling Via Nfkb
Hallmark Uv Response Dn
Hallmark Tgf Beta Signaling
Hallmark Apoptosis
Hallmark E2f Targets
Hallmark G2m Checkpoint
Hallmark Myc Targets V2
Hallmark Mtorc1 Signaling
Hallmark Uv Response Up
Hallmark Myc Targets V1
Hallmark Cholesterol Homeostasis
Δ AD1+AD2 vs pcDNA
p.adjust
0.05
0.04
0.03
0.02
0.01
0.00
GeneRatio
0.050
0.075
0.100
0.125
F
MSigDB Hallmark Gene Sets
Downregulated
Upregulated
Hallmark Myogenesis
Hallmark Epithelial Mesenchymal Transition
Hallmark Adipogenesis
Hallmark Protein Secretion
Hallmark Myc Targets V1
Hallmark E2f Targets
Hallmark Myc Targets V2
Hallmark G2m Checkpoint
Hallmark Mtorc1 Signaling
Hallmark DNA Repair
Δ TDU+AD1 vs pcDNA
p.adjust
0.05
0.04
0.03
0.02
0.01
0.00
GeneRatio
0.02
0.04
0.06
0.08
0.10

**Supplementary Figure S6: The hallmark myogenesis gene set is significantly downregulated in all the VGLL2::NCOA2 domain deleted mutant groups.**

(A-F) Hallmark gene sets enrichment plot detailing downregulated and upregulated gene sets in the full-length and all the domain-deleted VGLL2::NCOA2 fusion-oncogene transfected cells compared to the pcDNA3.1 control condition. Reads were aligned to a custom mouse genome (mm39) that also included the human VGLL2::NCOA2 fusion-oncogene. Key regulated pathways are indicated by asterisks and arrows.

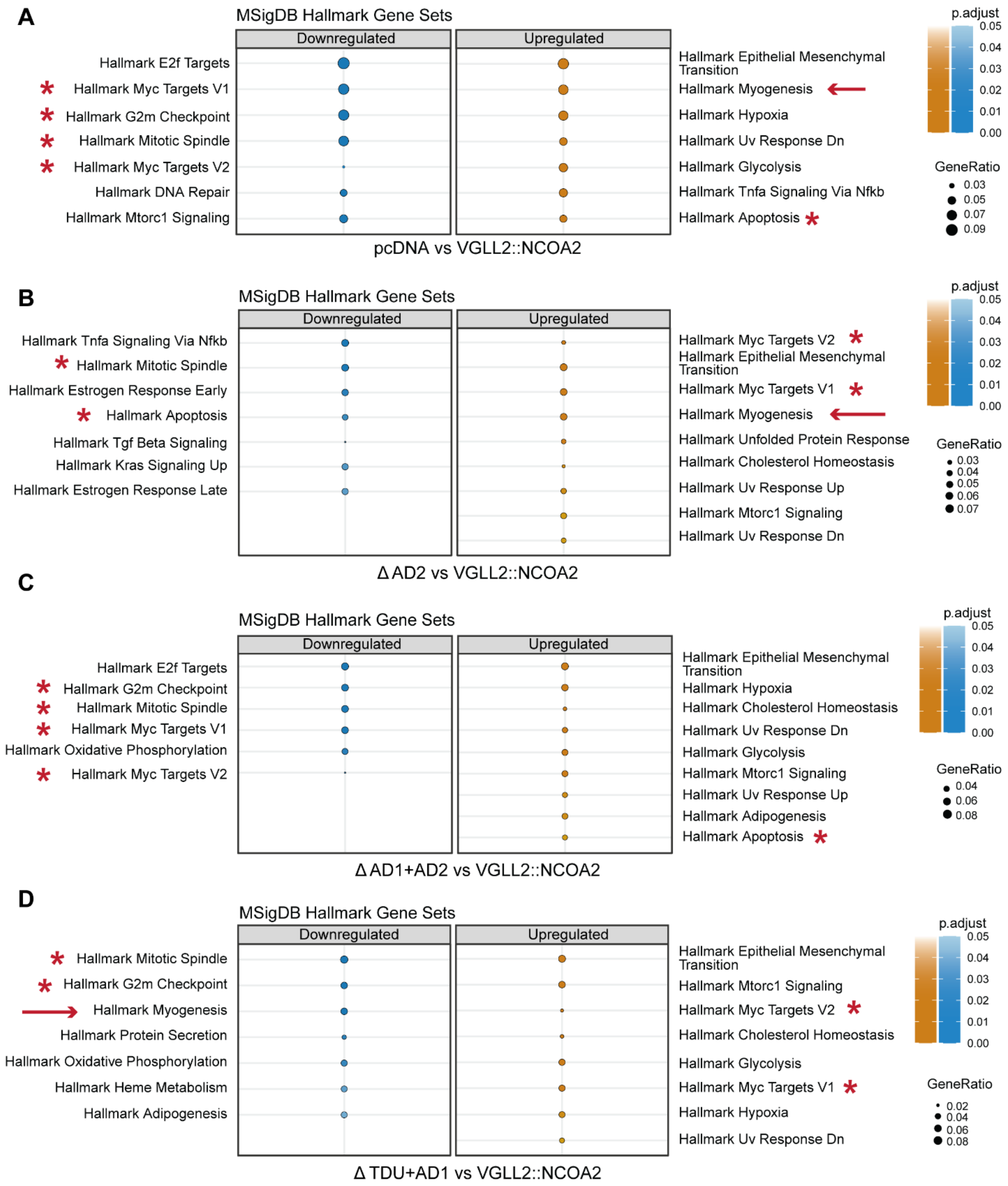


**Supplementary Figure S7: Transcriptomic comparison of VGLL2::NCOA2 AD2, AD1+AD2 and TDU+AD1 domain deletion mutants to the VGLL2::NCOA2 full-length condition.**

(A-D) Hallmark gene sets enrichment plot detailing downregulated and upregulated gene sets in pcDNA vector control, AD2, dual AD1+AD2, and dual TDU+AD1 deletion compared to the full-length VGLL2::NCOA2 condition. Reads were aligned to a custom mouse genome (mm39) that also included the human VGLL2::NCOA2 fusion-oncogene. Key regulated pathways are indicated by asterisks and arrows.

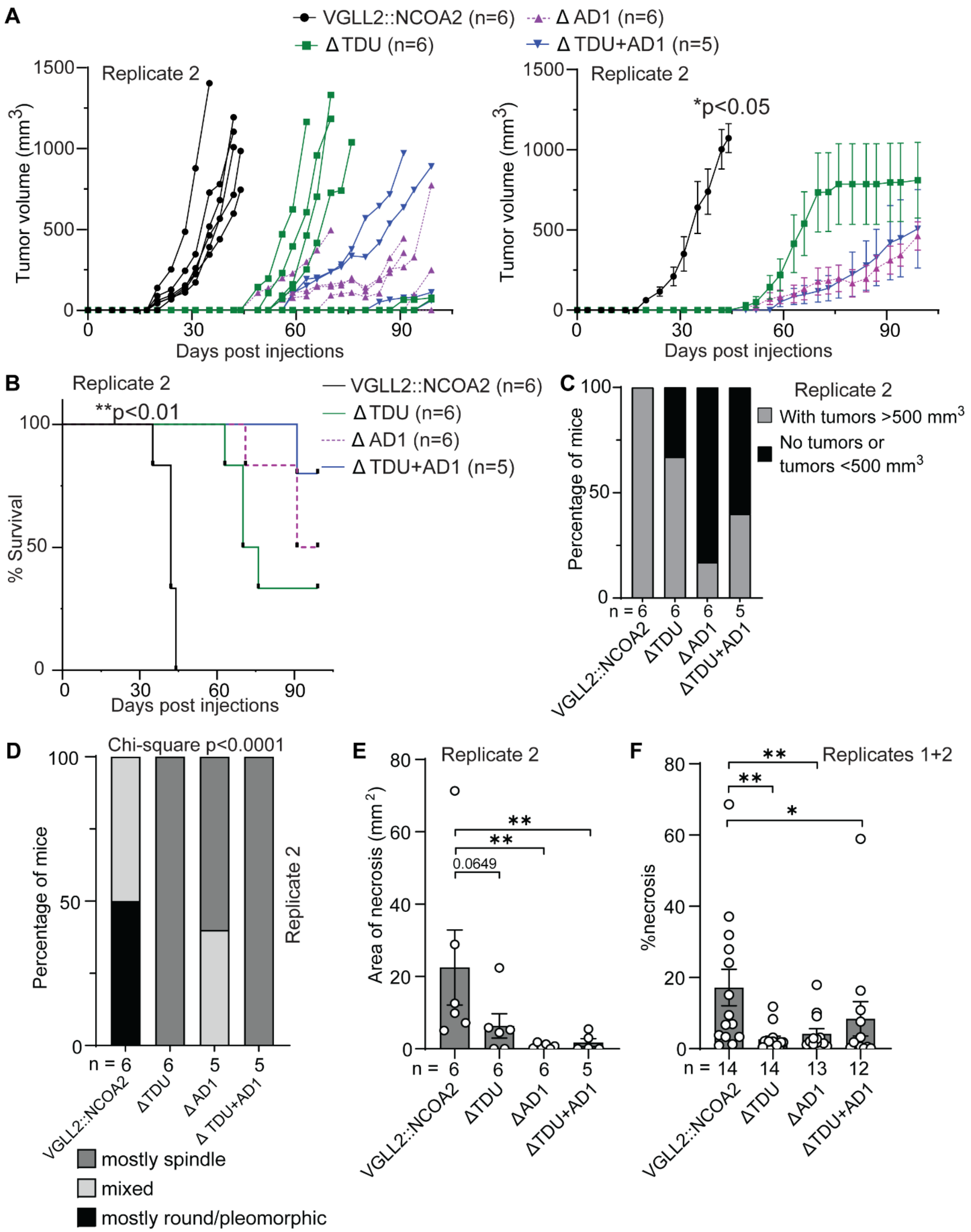

A
VGLL2::NCOA2 (n=6)
Δ AD1 (n=6)
Δ TDU (n=6)
Δ TDU+AD1 (n=5)
Replicate 2
Tumor volume (mm3)
Days post injections
*p<0.05
B
Replicate 2
**p<0.01
% Survival
Days post injections
C
Replicate 2
With tumors >500 mm3
No tumors or tumors <500 mm3
Percentage of mice
n = 6 6 6 5
VGLL2::NCOA2
ΔTDU
ΔAD1
ΔTDU+AD1
D
Chi-square p<0.0001
Percentage of mice
Replicate 2
n = 6 6 5 5
mostly spindle
mixed
mostly round/pleomorphic
E
Replicate 2
Area of necrosis (mm 2)
0.0649
**
n = 6 6 6 5
F
Replicates 1+2
%necrosis
*
n = 14 14 13 12

**Supplementary Figure S8: VGLL2::NCOA2 TDU and AD1 domains support fusion-driven tumor formation in a mouse allograft model – replicate experiment for Figure 5.**

(A) Tumor leg volume is plotted for each individual mouse as a function of days post injection for an independent replicate mouse allograft experiment from that presented in Figure 5 (left). Cumulative mean tumor volume of all the tumor-bearing mice as a function of days post injection (right). Data indicate mean ± standard error of mean (SEM). p values were calculated by comparing the mean days of tumor onset using a Kruskal Wallis non-parametric ANOVA test corrected for multiple comparisons using the 2-stage linear step-up procedure of Benjamini, Krieger, and Yekutieli. *$p<0.05$ with respect to comparing the full-length VGLL2::NCOA2 condition to all domain deletion mutant allograft tumors.

(B) Survival plot indicating the percentage of mice reaching tumor endpoint or humane endpoint for each group as a function of days post injection. Cumulative statistical analysis was performed using Log-rank (Mantel-Cox) test and individual tests between two groups were performed using Log-rank (Mantel-Cox) test and confirmed using Gehan-Breslow-Wilcoxon test. **$p<0.01$ with respect to comparing the full-length VGLL2::NCOA2 condition to all domain deletion mutant allograft tumors.

(C) Stacked bar plot indicating the number of mice with tumors greater than 500 $mm^3$ volume, and number of mice with no tumors and tumors smaller than 500 $mm^3$ volume.

(D) Pathological assessment of tumor cell morphology based on hematoxylin and eosin (HE)-stained slides in a semi-quantitative manner, using a three-tiered system: mostly (>75%) round/pleomorphic cells, mostly (>75%) spindle cells, or a combination of the above (mixed). Statistical significance was determined using a Chi-square test.

(E, F) Quantification of necrosis areas and percentage necrosis on hematoxylin and eosin (HE)-stained slides, was performed using the HALO image analysis platform and the HALO AI module (version 3.5.3577) from Indica Labs (Albuquerque, NM, USA). Data indicate mean ± standard error of mean (SEM). Statistical analysis comparing mean area of necrosis and mean %necrosis in the full-length VGLL2::NCOA2 group to the domain deletion mutants was performed using the Mann Whitney non-parametric test. *$p<0.05$, **$p<0.01$.

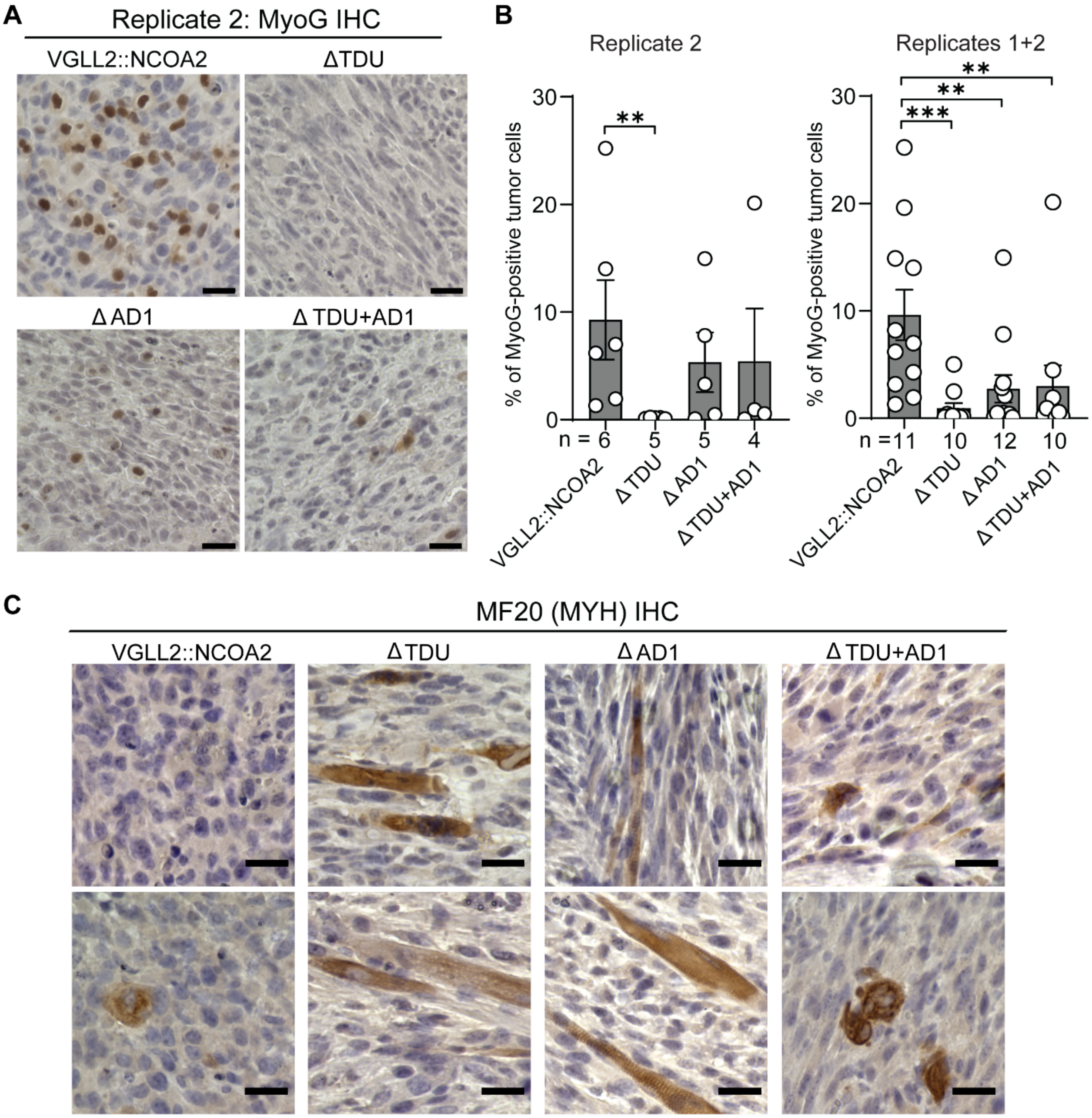


**Supplementary Figure S9: Expression patterns of myogenin and myosin-heavy chain in fusion-driven mouse allograft tumors.**

(A) Representative images of MyoG immunohistochemistry in the VGLL2::NCOA2, TDU deletion, AD1 deletion, and TDU and AD1 deletion conditions for allograft replicate 2. Scale bars: 25 µm.

(B) Quantification of the percentage of myogenin-positive tumor nuclei for allograft replicate 2 and combining replicates 1 and 2 on stained slides was performed using the HALO image analysis platform and the HALO AI module (version 3.5.3577) from Indica Labs (Albuquerque, NM, USA). Data indicate mean ± SEM. Statistical analysis comparing mean %myogenin-positive tumor cells in the mutant domain-deleted fusion-oncogene group to the full-length fusion-oncogene group was performed using the Mann Whitney non-parametric test. **$p<0.01$, ***$p<0.001$.

(C) Representative images of all myosin heavy chains (MYH) immunohistochemistry (MF20) in the VGLL2::NCOA2, TDU deletion, AD1 deletion, and TDU and AD1 deletion conditions. Scale bars: 25 μm.
